# Predicting the efficacy of BH3 mimetics through the profiling of multiple protein complexes in acute myeloid leukemia

**DOI:** 10.1101/2023.01.16.524337

**Authors:** Changju Chun, Ja Min Byun, Minkwon Cha, Hongwon Lee, Byungsan Choi, Hyunwoo Kim, Saem Hong, Yunseo Lee, Hayoung Park, Youngil Koh, Tae-Young Yoon

## Abstract

BH3 mimetics are protein-protein interaction (PPI) inhibitors that saturate anti-apoptotic proteins in the BCL2 family to induce apoptosis in cancer cells, a prominent example of protein complex-targeting therapies. Despite the remarkable success of the BH3 mimetic ABT-199 in treating hematological malignancies, only a fraction of patients respond to ABT-199 and eventually develop resistance, necessitating the predictive biomarkers for both initial responses and resistance development. We here used the single-molecule pull-down and co-immunoprecipitation platform to quantify more than 20 different types of PPI complexes using ∼1.2×10^6^ cells in total, revealing the rewired status of the BCL2 family PPI network. By comparing the obtained multi-dimensional data with BH3 mimetic efficacies determined *ex vivo*, we constructed an analysis model for ABT-199 efficacy that designates the BCL2-BAX and BCLxL-BAK complexes as the primary mediators of drug effectiveness and resistance. We then applied this model to assist in therapeutic decision-making for acute myeloid leukemia patients in a prospective manner. Our work demonstrates a capability for the extensive characterization of PPI complexes in clinical specimens, potentially opening a new avenue of precision medicine for protein complex-targeting therapies.

---

Cells employ large protein complexes to make intricate decisions, reflecting diverse cellular states^1,2^. Prominent examples include regulation by mammalian target of rapamycin (mTOR) complexes in metabolism^3^, inflammasomes in innate immunity^4^ and BCL2 family complexes in apoptotic cell death^5^. Despite the unexpected breadth of associated proteins and their rapid turnover in these complexes, connectivity amongst several core components still defines critical setpoints in cellular decision-making. For example, across possible combinations of the BCL2 family proteins, the formation of proteinaceous pores by BAX and BAK leads to mitochondria outer membrane permeabilization (MOMP), making a point of no return for apoptosis initiation^6–8^. The survival and apoptotic pressure accrued in cells are reflected by the expression of anti- and pro-apoptotic proteins, tipping the scale toward survival or apoptosis, respectively. While BCL2, BCLxL, and MCL1 make up the group of anti-apoptotic proteins that sequester BAX/BAK to prevent pore formation^5,9–11^, pro-apoptotic proteins (such as BIM, BAD, and BID) share an alpha-helical BH3-domain and interact with either anti-apoptotic proteins or BAX/BAK^12–14^, thereby unleashing the activation of BAX/BAK for pore formation^15,16^.

In diseased states, cellular decisions are often misguided by chronic or spurious formation of these protein complexes, as seen with mTOR complexes found in various cancers or type 2 diabetes^17,18^ and inflammasomes in irritable bowel syndrome or neurodegenerative diseases^19,20^. Likewise, many cancers exhibit upregulated PPIs of the anti-apoptotic proteins to suppress the activity of BAX/BAK, thus averting apoptosis initiation^10,16,21–23^. Consequently, extensive efforts have been made to develop PPI inhibitors that can disintegrate disease-causing protein complexes^24^. In particular, BH3 mimetics, small-molecule drugs that structurally emulate the BH3 domain, are a notable example in these endeavors^25,26^. While sparing pro-apoptotic proteins, BH3 mimetics are expected to bind selectively to the binding grooves of the anti-apoptotic proteins to shut down their interactions with pro-apoptotic proteins and BAX/BAK^27^. By introducing an azaindole moiety to ABT-263 (Navitoclax) that binds to both BCL2 and BCLxL^28^, ABT-199 (Venetoclax) has been developed to specifically bind to BCL2 with a sub-nM affinity^29^. ABT-199 was approved for the treatment of chronic lymphoid leukemia (CLL) in 2016 (NCT01889186)^30^ and acute myeloid leukemia (AML) in 2018 (NCT02287233, NCT02203773)^31,32^, which stimulated the development of other BH3 mimetics that target other anti-apoptotic proteins beyond BCL2^27,33^.

In line with other targeted therapies, a precision medicine platform could facilitate the customized use of PPI inhibitors, using the unique molecular information of individual patients^34^. Despite ABT-199’s remarkable success in treating CLL^35,36^, over 30% of AML patients still fail to respond^32,37,38^, and many with initial favorable responses eventually develop resistance^39,40^. These facts underscore unmet needs where diverse treatment options could benefit patients if initial responses or resistance development could be accurately predicted. However, multiple factors, including post-translational modifications, protein conformational changes, and cellular localizations, may hinder the direct translation of genomic and proteomic information into accurate PPI strength predictions^5,41,42^. Genomic and proteomic profiling have thus found limited use in this technical niche^43–45^, and correlations between total expression levels of BCL2 and other anti-apoptotic proteins and response to BH3 mimetics have not been robust enough for clinical application^46–49^. While the BH3 profiling method offers potential usefulness in predicting BH3 mimetics, it is not routinely used in clinical settings, likely due to the need for sample viability maintenance^50–52^.

In this work, we hypothesize that the responsiveness of individual cancers to a PPI inhibitor would be determined by the ever-evolving compositions of the target protein complexes. Specifically, in the case of BH3 mimetics, the heterocomplexes between anti- and pro-apoptotic proteins and their intricate balances may primarily mediate the efficacy of BH3 mimetics in given cancers. We propose that a predictive algorithm may be trained by extensively profiling BCL2 family protein complexes in as many cancers with known responses as possible. To enable such profiling with minimal sample consumption and increased throughput, we adopted the single-molecule pull-down and co-immunoprecipitation (SMPC) platform, directly immobilizing target protein complexes on the imaging plane of a single-molecule fluorescence microscope and counting the number of complexes^53–56^. This approach allowed us to determine the populations of 20 different types of protein complexes (and protein levels) using only ∼30,000 cells per complex type, revealing the latest rewiring status of the BCL2 family network in individual clinical samples.

We found that the introduction of bacterial toxin and BH3 mimetics caused significant changes in the composition of BCL2 protein complexes, with only minor alterations in corresponding protein levels. This finding underscores the importance of directly profiling protein complexes rather than merely analyzing protein levels. By correlating these multi-dimensional data with the *ex vivo* determined efficacies of BH3 mimetics, we were able to train analytical models that identified the protein and PPI parameters playing pivotal roles in mediating the efficacy of BH3 mimetics. For ABT-199, we found that the counts of two specific complexes, BCL2-BAX and BCLxL-BAK, served as critical contrasting analysis parameters, with the former driving ABT-199’s efficacy and the latter contributing to resistance development. Following a similar procedure, we constructed an efficacy model for an MCL1-targeting inhibitor. Subsequently, we applied the developed analytical model to guide therapeutic decisions for AML patients in a prospective fashion, confirming the model’s predictive power. Our work showcases the utility of the SMPC platform for extensive and accurate profiling of protein complexes and presents a path toward precision medicine in therapies targeting protein complexes.

## Results

### Profiling BCL2 family protein complexes using SMPC

To assess BCL2 family protein complexes and gain insights into how apoptosis is suppressed in individual cancers, we utilized the SMPC platform to characterize the PPI network among BCL2 family proteins^53,54,56,57^. We initiated surface immunoprecipitation (IP) of one of the primary anti-apoptotic proteins (i.e., BCL2, BCLxL, or MCL1) from crude cell extracts (Fig. 1a and Extended Data Fig. 1a-c), followed by the addition of a monoclonal antibody that binds to the interaction partner complexed with the surface-immobilized bait. A fluorescently labeled secondary antibody was then used, completing an immunoassay to detect specific protein complexes (CPXs) (Fig. 1b, left). Additionally, we employed a detection antibody that binds to an epitope directly on the surface bait to determine the total amount of surface-immobilized baits (protein total level, LV) (Fig. 1b, right and Extended Data Fig. 1d-f).

**Fig. 1.**
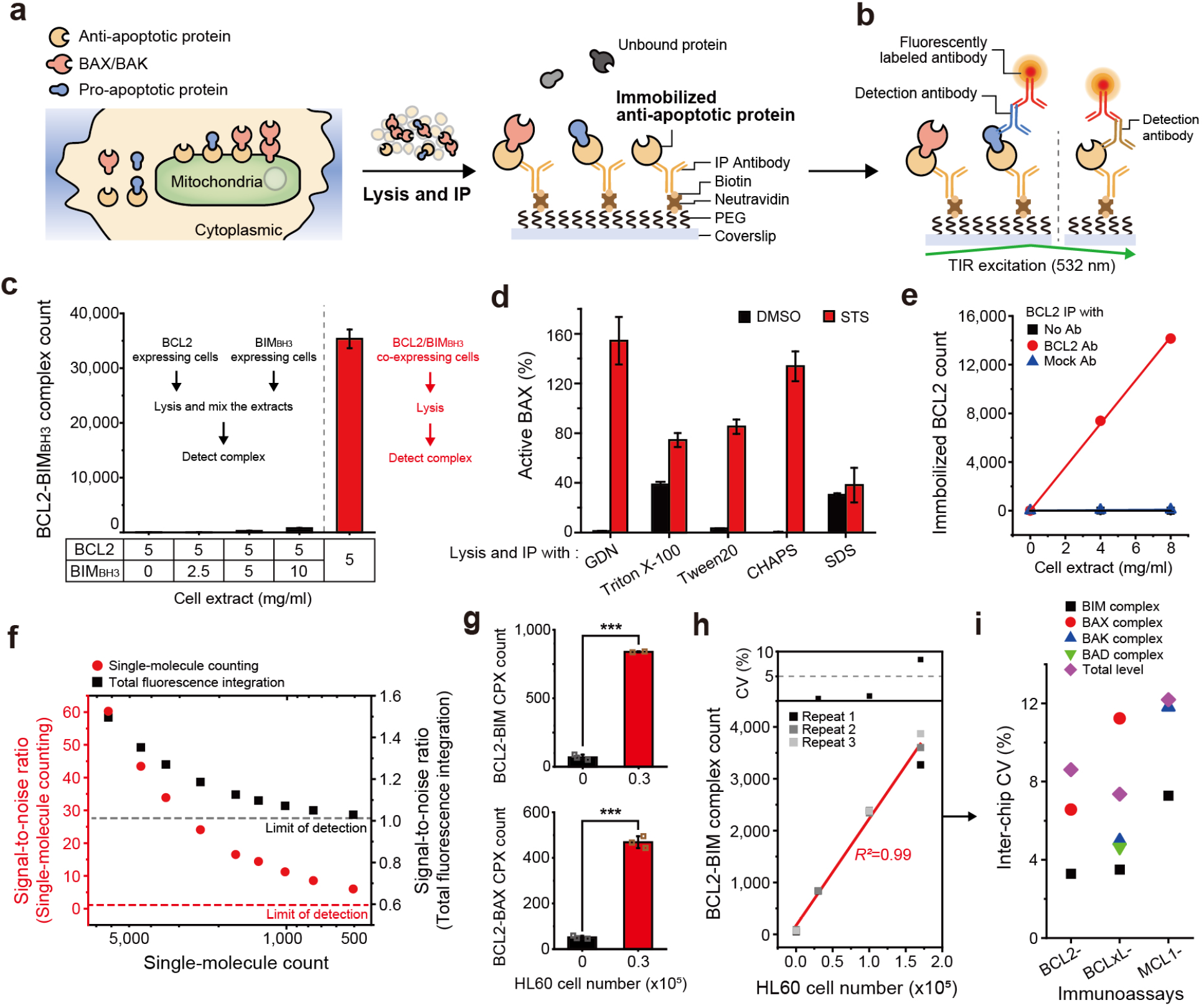
| Profiling BCL2 family protein complexes using the SMPC platform. **a**, Schematic for the surface IP of BCL2 family proteins. Anti-apoptotic proteins (BCL2, BCLxL, MCL1) in crude cell extracts were immobilized onto the surface of the reaction chamber. **b**, Schematic of immunoassay selectively detecting protein complexes (left) and the total level of surface bait protein (right). The fluorescence signals from detection antibodies were imaged under total internal reflection (TIR) excitation. **c**, Comparing the formation of model BCL2-BIM_BH3_ complex in an intracellular environment by BCL2-mCherry/BIM_BH3_-eGFP co-transfection (red) and *in vitro* mixing of two individual extracts with each expressing BCL2-mCherry and BIM_BH3_-eGFP (black). **d**, Relative changes of active BAX level measured for STS treated HL60 cells after lysed with different detergent types. The monoclonal antibody 6A7 was used to detect the activated BAX^60^. **e**, Surface-immobilization of BCL2-eGFP from the crude extract with indicated IP antibodies. **f**, Comparison of signal-to-noise ratio between single-molecule counting analysis (red) and integration of total fluorescence signals (black). **g**, Counts of endogenous BCL2 protein complexes (BCL2-BIM, BCL2-BAX) from the 30,000 HL60 cells by using immunoassay. Error bars represent mean±s.d from independent inter-chip measurement (*n*=3). (Two-sided two-sample *t*-test, *p*-value=1.5e-5, 1.1e-5 respectively). **h**, Counts of endogenous BCL2-BIM complexes and inter-chip CVs from the fixed numbers of HL60 cells (*n*=3). **i**, Inter-chip CVs obtained from independent inter-chip measurement for all the immunoassays and cell numbers (*n*=3). Error bars represent means±s.d.

To enhance sensitivity and reproducibility in this protein complex detection method, we optimized the lysis protocol^57,58^. By using HEK293T cells co-expressing BCL2 and the BH3 domain of BIM (BIM_BH3_), we found that mild lysis with cholesterol-like detergents, such as glycol-diosgenin (GDN) and digitonin, minimized dissociation of the model BCL2-BIM complexes (Extended Data Fig. 1g). Furthermore, this lysis protocol did not induce the formation of spurious complexes even in dense cell extracts (i.e., at a total protein concentration of 5 to 10 mg/ml), confirming that all detected protein complexes were formed in cellular environments (Fig. 1c). Conformational changes of the BCL2 family proteins during their activation are crucial in their function^16,59^. For instance, the pore formers, BAX and BAK, expose their α1 helices in the activated state, which are otherwise buried in the core structure. By assessing the active status of BAX using an anti-α1 helix antibody, we found that the detergent condition significantly affected BAX’s conformational state^2,60^. Triton X-100, often used in conventional co-IP methods, induced spurious conformational changes^61^, while GDN essentially preserved BAX’s active conformations induced by the addition of a bacterial toxin, staurosporine (STS)^62^ (Fig. 1d). Similarly, we experimentally confirmed the ability to assess the activation status of BAK^63,64^, by employing an antibody that specifically binds to the α1 helix of BAK (Extended Data Fig. 1h,i). Finally, we experimentally confirmed that a single freeze-thaw cycle did not seriously affect the single-molecule counts of BCL2 protein complexes (Extended Data Fig. 1j-l).

To minimize non-specific adsorption to the surface, we placed a high-density polyethylene glycol (PEG) layer between the IP antibodies and the glass coverslip^57,58^ (Fig. 1a). This PEG layer proved essential in enabling direct surface IP of the target protein complexes from the crude extracts reducing background signals to a minimal level on our single-molecule fluorescence imaging (Fig. 1e). By effectively removing background signals through signal processing, we could identify single-molecule fluorescence spots and directly count their numbers (Extended Data Fig. 2a-c)^55,65^. Compared with conventional approaches that integrate total fluorescence signals over a field of view, this single-molecule counting method maintained signal-to-noise ratios higher than 5 when there were ∼1,000 or fewer individual single-molecule spots per imaging area (Fig. 1f and Extended Data Fig. 2).

Through the combination of the three technological factors of optimized lysis protocol, anti-fouling PEG layer, and direct single-molecule counting, we achieved statistically significant counts for endogenous BCL2-BIM or BCL2-BAX CPXs using only ∼30,000 AML cells (Fig. 1g,h). We also demonstrated a similar capability using a non-small cell lung cancer cell line (PC9 cells), where we detected 8 different BCLxL- and MCL1-protein complexes using less than 50,000 PC9 cells each (Fig. 1i and Extended Data Fig. 3). Three independent inter-chip measurements yielded coefficients of variation (CVs) of less than 15% for all assays and cell numbers tested, confirming the high stability of the established SMPC platform (Fig. 1h,i and Extended Data Fig. 3).

### Tracking the responses to apoptotic stress using SMPC

To investigate if the developed SMPC platform could monitor alterations within the BCL2 family PPI network, HL60 cells were treated with 2 μM of STS. The crude extracts were obtained from these cells at various time points during the treatment (Extended Data Fig. 4a). Through flow cytometry analysis, the early apoptosis population, characterized by high Annexin V staining, was observed to surge at 1 hour of STS treatment, reducing the healthy cell population (Extended Data Fig. 4a,b). The late apoptosis population, stained with both Annexin V and PI, increased after 4 hours. Necrosis populations remained negative throughout the study, signifying that the cells underwent apoptosis rather than necrosis.

We observed that the counts of active BAX and BAK dramatically increased as apoptosis progressed (Fig. 2a). The count of BAX-BAK heterocomplexes, a more direct surrogate marker of MOMP^66,67^, began to rise simultaneously with the active BAX and BAK counts, but continued to escalate into the late apoptosis stage. We then quantified the total levels of anti-apoptotic proteins (Fig. 2b). Through careful calibration of immunolabeling efficiencies (Extended Data Fig. 5a-c), we were able to directly compare the quantities of the primary anti-apoptotic proteins and their protein complexes (Fig. 2b-d). We found that BCL2 proteins were more abundant by 4 to 5 times compared to BCLxL and MCL1, aligning with the known dependence of HL60 on BCL2 (Fig. 2b). As apoptosis progressed, BCLxL and MCL1 levels gradually diminished, while BCL2 protein levels exhibited a slight yet significant increase.

**Fig. 2.**
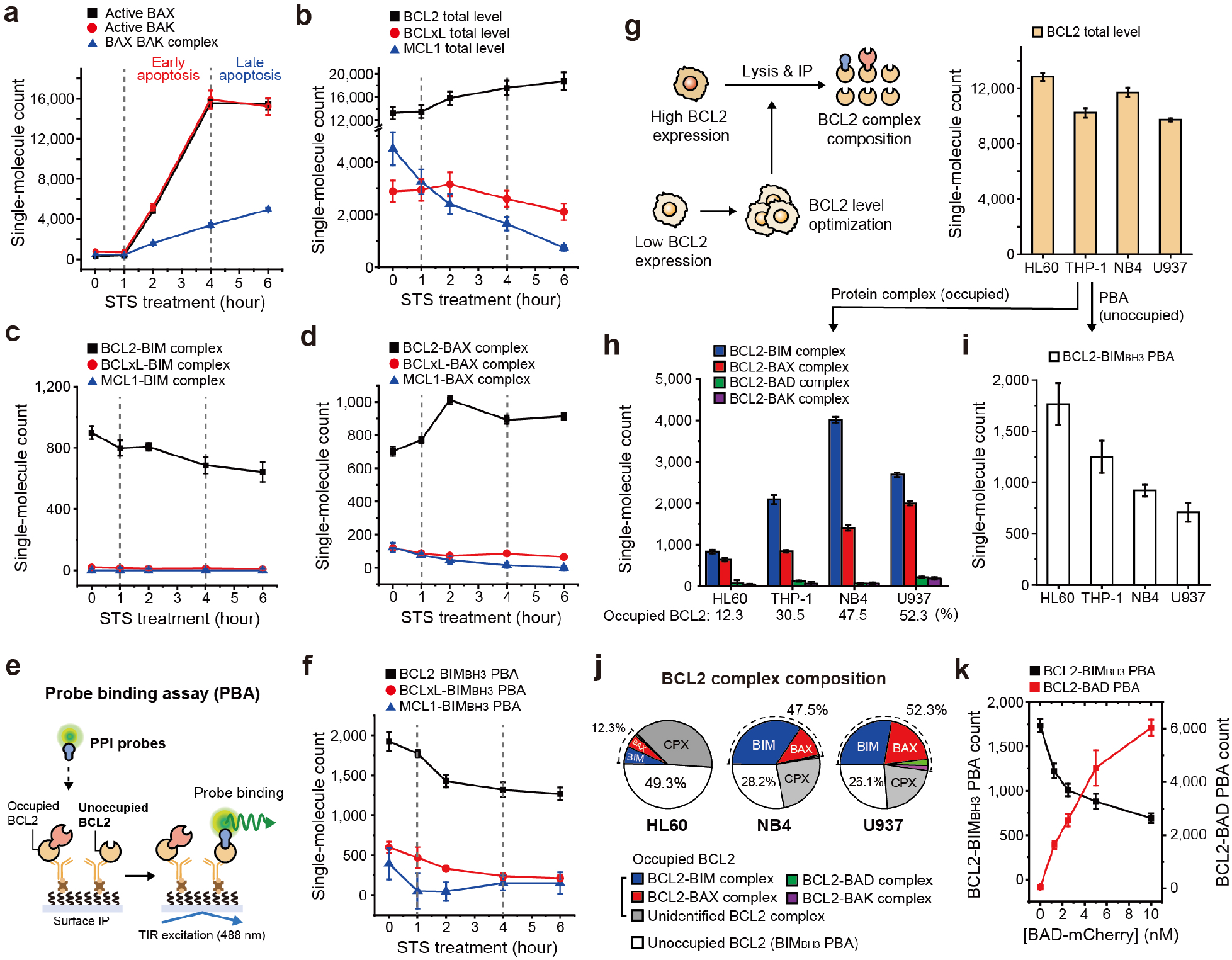
| Tracking the responses to intrinsic apoptotic stress with SMPC. **a-d**, Changes of endogenous BCL2 family PPI profiles of HL60 cells through the apoptosis progression by 2 μM of STS. (**a**) Active BAX/BAK level, (**b**) Total levels of anti-apoptotic proteins (BCL2, BCLxL, MCL1), (**c**) BIM complexes with anti-apoptotic proteins, (**d**) BAX complexes with anti-apoptotic proteins. **e**, Schematic for the probe binding assay (PBA) to measure the unoccupied populations of surface-immobilized anti-apoptotic proteins. **f**, Changes of BIM_BH3_ PBAs for anti-apoptotic proteins from HL60 cells through the apoptosis progression by STS. **g**, Schematic for the comparison of the BCL2 protein complex compositions in different AML cell lines. The density of surface-immobilized BCL2 was constantly maintained by optimizing the total protein concentration of crude cell extracts for all AML cell lines (HL60, THP-1, NB4, U937). **h**, Compositions of the BCL2 complexes (BIM, BAX, BAD, BAK) in four different AML cell lines. **i**, BCL2-BIM_BH3_ PBA counts of four AML cell lines. **j**, The BCL2 complex compositions in HL60, NB4, and U937 cell extracts. **k**, Changes of BCL2-BIM_BH3_ PBA and BCL2-BAD PBA with increasing amounts of BAD probe. BIM_BH3_ probe was presented in 10 nM. Error bars represent means±s.d. The single-molecule counts were rescaled to account for the labeling efficiencies of the immunoassays (Extended Data Fig. 5) as well as the specific incubation conditions.

Contrary to the minimal change in BCL2 protein level, the heterocomplexes formed between BCL2 and pro-apoptotic proteins displayed dynamic changes throughout the apoptosis stages. We discovered that BCL2 protein complexes outnumbered those involving BCLxL and MCL1, further confirming HL60’s BCL2-dependence (Fig. 2c,d). Notably, the two protein complexes exhibited contrasting behaviours. While the counts of the BCL2-BIM complex gradually decreased over time, the BCL2-BAX complex counts surged for the first 2 hours following STS treatment (Fig. 2c,d). Collectively, our comprehensive profiling of the BCL2 family protein complexes revealed that BCL2 proteins mainly functioned to sequester the active BAX proteins during STS-induced apoptotic pressure in HL60, while simultaneously releasing some previously held BIM proteins. Presumably, under the applied STS treatment conditions, the active BAX outnumbered the BCL2’s sequestration capacity, leading to the HL60 cells succumbing to apoptotic stress and initiating MOMP (Fig. 2a,d).

### Assessment of potential PPIs using protein interactor probes

Our findings reinforce the understanding that BCL2 family proteins, akin to many cell signalling proteins, function by binding to (or sequestering) interaction partners^4^. During elevated survival pressure in HL60 cells induced by STS treatment, the BCL2 proteins consistently formed additional BCL2-BAX complexes to counteract apoptotic pressure (Fig. 2d). This led us to conceive additional assays to evaluate the capacity of BCL2 family proteins to form further protein complexes beyond current complex populations.

To evaluate the affinities of potential PPIs, we employed the same surface pull-down approach to immobilize anti-apoptotic BCL2 family proteins, a procedure confirmed to cause minimal IP crosstalk and non-specific adsorption (Fig. 2e). Various pro-apoptotic BCL2 proteins, tagged with enhanced green fluorescent proteins (eGFPs), were then introduced into the reaction chamber. We hypothesized that these PPI probes would engage with a pool of anti-apoptotic proteins that were unoccupied but still capable of forming potential protein complexes (Fig. 2e). Indeed, this PPI probe binding assay not only allowed us to replicate the dissociation constants for the binding interactions between the anti- and pro-apoptotic proteins, but also faithfully reproduce known biochemical features (Extended Data Fig. 6a-h)^5,52^. These included a higher affinity of BCL2 for BAD compared to BIM and negligible interactions for BCL2-NOXA and MCL1-BAD pairs (Extended Data Fig. 6e,f)^4^. We further verified that these PPI probes did not disrupt the pre-existing, endogenous protein complexes (Extended Data Fig. 6i).

Utilizing this probe binding assay (PBA), we analyzed BCL2 proteins extracted from the STS-treated HL60 cells (Extended Data Fig. 4b). By employing the BH3 domain of BIM (BIM_BH3_-eGFP) as a PPI probe, BCL2 exhibited a significantly higher PBA count in comparison to BCLxL and MCL1 (Fig. 2f). Given the higher affinities of the BIM_BH3_ domain for BCLxL and MCL1 than for BCL2 (Extended Data Fig. 6b), this elevated PBA count suggests that BCL2 works as a larger reservoir in sequestering pro-apoptotic proteins compared to BCLxL or MCL1 in HL60 (Fig. 2f). Strikingly, as apoptosis progressed, the PBA count of BIM_BH3_ declined, despite the rise in total BCL2 levels, possibly indicating the exhaustion of available BCL2 proteins through the formation of BCL2-BAX complexes (Fig. 2c-f). Furthermore, we inspected the composition of the BCL2 protein complexes in four AML cell lines (Fig. 2g). When immobilizing an equivalent number of total BCL2 proteins per view (Fig. 2g, right), NB4 and U937 cells were found to contain substantially more BCL2-BIM and BCL2-BAX complexes, the two main species of BCL2 protein complex species, compared to HL60 (Fig. 2h). This observation corresponded with the PBA count being more than halved for NB4 and U937 relative to HL60 (Fig. 2i), reflecting increased BCL2 occupancy and thereby reduced pools of BCL2 proteins available for potential PPIs in these cells (Fig. 2j). It was noteworthy that the combined totals of the PBA and protein complex counts accounted for 60 to 80% of the total BCL2 proteins in the cell lines we investigated. This suggests that a significant proportion of BCL2 proteins may be complexed with interaction partners not accounted for in our study (Fig. 2j).

Finally, we investigated whether different PPI probes target the same pool of BCL2 proteins or distinct, separate BCL2 pools (i.e., each pool accessible by only specific PPI probes). To answer this question, we introduced multiple PPI probes simultaneously and found that these probes directly competed with one another, as well as with ABT-199 (Fig. 2k and Extended Data Fig. 6j,k). In addition, the consistency of the PBA counts across different cell lines and PPI probes indicated the existence of a singular, common BCL2 protein pool, rather than multiple, fragmented pools of BCL2 proteins (compare Fig. 2i with Extended Data Fig. 6l-n).

### Tracing changes in the BCL2 family protein complexes provoked by BH3 mimetics

Leveraging the two assays we established (namely, single-molecule immunoassay and PBA), we set out to explore how BH3 mimetics instigated apoptotic pressure in AMLs. We exposed HL60 cells to 100 nM ABT-199 for 24 hours, resulting in a modest ∼30% increase in the total BCL2 level, while BCLxL and MCL1 levels remained minimal (Fig. 3a). Conversely, the amounts of active BAX and BAK, along with BAX-BAK heterocomplexes, surged five-fold after ABT-199 treatment, signalling active apoptosis induction in HL60 (Fig. 3b). Although total BAX and BAK populations were largely consistent (Fig. 3c), the majority of the BCL2-BIM and BCL2-BAX complexes dissociated due to ABT-199 exposure (Fig. 3d). These findings highlight that BAX, once released from BCL2, mainly contributed to the steep increases in active BAX and BAK populations. Additionally, ABT-199’s liberation of BCL2 proteins led to an elevated PBA count – a direct contrast to the STS-treated case where the PBA count dwindled over the treatment period (Fig. 3e).

**Fig. 3.**
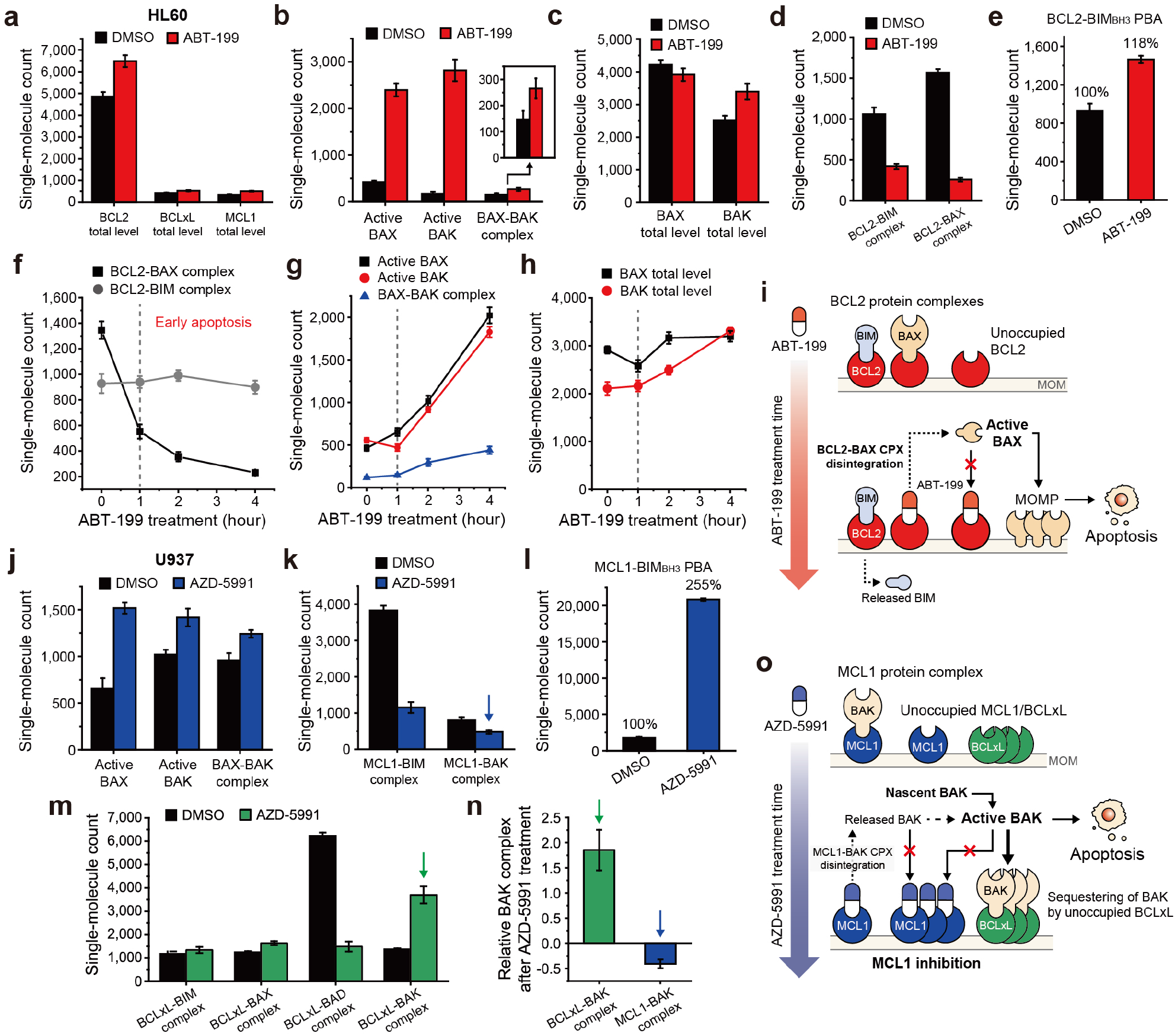
| Changes in the BCL2 family protein complexes provoked by BH3 mimetics. **a-e**, Changes of BCL2 family PPI profile in HL60 cells after ABT-199 treatment (100 nM, 24 hours). (**a**) Total levels of anti-apoptotic proteins, (**b**) Active BAX/BAK level, (**c**) Total levels of BAX/BAK, (**d**) BCL2 complexes (BIM, BAX), (**e**) BCL2-BIM_BH3_ PBA. **f-h**, Changes of BCL2 family PPI profile in HL60 cells after higher concentration of ABT-199 treatment (300 nM, 4 hours). (**f**) BCL2 complexes (BIM, BAX), (**g**) Active BAX/BAK level, (**h**) Total levels of BAX/BAK. **i**, Schematic illustration of the mode of action of ABT-199. **j-l**, Changes of BCL2 family PPI profile in U937 cells after AZD-5991 treatment (100 nM, 24 hours). (**j**) Active BAX/BAK level, (**k**) MCL1 complexes (BIM, BAK), (**l**) MCL1-BIM_BH3_ PBA. The decrease of MCL1-BAK complex after AZD-5991 treatment was indicated (blue). **m**, Changes of BCLxL complexes (BIM, BAX, BAD, BAK) from U937 cells after AZD-5991 (100 nM, 24 hours) treatment. The increase of BCLxL-BAK complex after AZD-5991 treatment was indicated (green). **n**, Comparison of relative changes of BAK complexes from U937 cells after AZD-5991 treatment. The relative changes were obtained from the indicated data in (**k**) and (**m**). **o**, Schematic illustration of the mode of action of AZD-5991. (**d**,**k**,**m**) The single-molecule counts were rescaled to account for the labeling efficiencies of the immunoassays. Error bars represent means±s.d.

To delineate the dynamics of BCL2-BIM and BCL2-BAX complexes, especially at early ABT-199 treatment stages, we administered a higher 300 nM concentration of ABT-199 to HL60 cells and conducted PPI profiling at each hourly interval following initial treatment (Extended Data Fig. 4c). The two major BCL2 complexes showed opposing results. The BCL2-BAX complex began to unravel instantly after ABT-199 administration, while the number of BCL2-BIM complexes largely persisted (Fig. 3f). The kinetics of BCL2-BAX dissociation synchronized with the activation of BAX/BAK conformations and BAX-BAK heterocomplexes (Fig. 3g). This data suggested that the dissociation of BCL2-BAX complexes – without necessitating BCL2-BIM dissociation or additional BAX expression – likely sufficed to invoke pro-apoptotic pressure and initiate apoptosis (Fig. 3f-i)^43,68–70^.

We further investigated whether similar complex dynamics were observable in MCL1 inhibitor-responsive cells. In reaction to AZD-5591^71^, U937 cells exhibited augmented populations of active BAX and BAK proteins, along with BAX-BAK heterocomplexes, consistent with U937’s susceptibility to the MCL1 inhibitor (Fig. 3j). Mirroring the case with ABT-199-treated HL60 cells, AZD-5591 treatment spurred a conspicuous dissociation of MCL1 protein complexes (MCL1-BIM and MCL1-BAK), evidenced by the pronounced increase in BIM_BH3_ probe binding for MCL1 (Fig. 3k,l). However, unlike the HL60 case where other anti-apoptotic proteins demonstrated minimal alterations, BCLxL proteins in U937 cells exhibited significant surges in their complex counts, with BCLxL-BAK complexes becoming preeminent (Fig. 3m). A thorough calibration of labeling efficiencies allowed us to discern that the observed rise in BCLxL-BAK markedly outpaced the decrease in MCL1-BAK complexes (Fig. 3n and Extended Data Fig. 5a,c). We consequently hypothesized that the majority of new BCLxL-BAK complexes originated from the sequestering of nascent BAK proteins, rather than capture from those liberated from MCL1 (Fig. 3o). In summation, these data substantiate the feasibility of comprehensive profiling of BCL2 family protein complexes, enhancing our understanding of how apoptotic pressure is both generated and managed by the BCL2 family protein network within individual cancers.

### Constructing an analysis model for ABT-199 efficacy with multi-dimensional PPI profiling

So far, our collective data suggest that BH3 mimetics work through either protein complex disintegration, saturation of anti-apoptotic proteins, or both to achieve their effects. Additionally, preliminary findings hint at a single-agent response when cancer cells show a specific dependence on a singular anti-apoptotic protein, with minimal compensation from other anti-apoptotic proteins (Extended Data Fig. 7). We therefore decided to investigate whether multi-parameter data could be used to develop an analytical model to identify the dependency on anti-apoptotic proteins for specific cancers and forecast their responsiveness to BH3 mimetic drugs.

For this purpose, we analyzed 32 primary AML samples, of which 30 were gathered from bone marrow mononuclear cells (BMMCs) and 2 from peripheral blood mononuclear cells (PBMCs) (Fig. 4a). The clinical diagnoses included *de novo* AML (*n*=17), relapsed AML (*n*=4), and secondary AML (*n*=11) (Supplementary Table 1). We evaluated the *ex vivo* effectiveness of ABT-199, determining a normalized area under the response curve (AUC) for each specimen (Fig. 4a, upper). Concurrently, the SMPC PPI profiling was performed on the same cohort samples in a blind fashion, allowing the determination of 22 unique protein/PPI metrics for each sample comprised of roughly 1.2×10^6^ cells in total (Fig. 4a, lower and Supplementary Table 2).

**Fig. 4.**
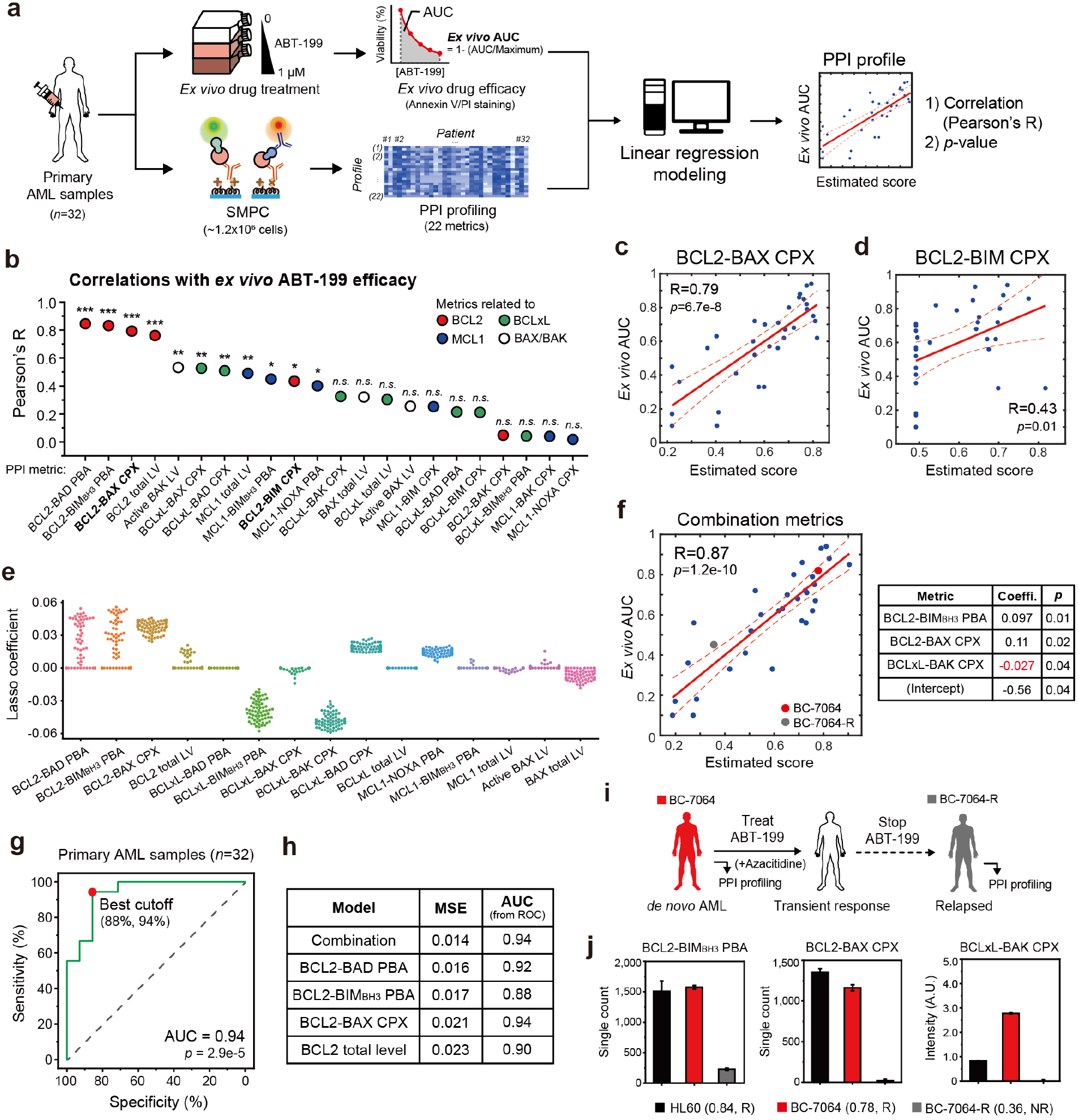
| Constructing an analysis model for ABT-199 efficacy with multi-dimensional PPI profiling. **a**, Schematic for generating linear regression correlation between *ex vivo* efficacy of ABT-199 and PPI profiles from primary AML samples. The collected AML cells were cultured and treated with 0-1 μM of ABT-199, and the area under curves (AUCs) of cell viability were obtained as *ex vivo* drug efficacy (upper). ∼1.2×10^6^ of primary AML cells on the same cohort underwent PPI profiling with the SMPC platform (lower) (*n*=32). **b**, Pearson correlations between *ex vivo* AUC of ABT-199 and PPI metrics for primary AML samples (*p-value*: *<0.05, **<0.01, ***<0.001, *n.s.*=not significant). **c**,**d**, Correlations between *ex vivo* AUC and single BCL2-related PPI metrics. (**c**) BCL2-BAX CPX, (**d**) BCL2-BIM CPX. **e**, Lasso coefficients of PPI profiles correlated with *ex vivo* AUC of ABT-199 for primary AML samples (67 models). **f**, Correlation between *ex vivo* AUC and combination of multiple PPI metrics (BCL2-BIM_BH3_ PBA, BCL2-BAX CPX, and BCLxL-BAK CPX). The statistical indicators (coefficient and *p*-value) of each metric were presented. **g,** Receiver operating characteristic (ROC) curve between the estimated score and the *ex vivo* efficacy. **h,** Comparison of predictive powers between analysis models. **i**, Clinical features and the ABT-199 administration history of BC-7064. **j**, Comparison of the PPI profiles and the estimated scores with PPI diagnostic results (R: responsive, NR: non-responsive) from the initial and the relapsed BC-7064 samples. Error bars represent means ± s.d.

Upon comparing the protein/PPI assay data with AUC, four BCL2-related metrics – BCL2-BAD PBA, BCL2-BIM_BH3_ PBA, BCL2-BAX CPX and BCL2 total level – exhibited the highest correlations with *ex vivo* efficacy (Fig. 4b). Although BCLxL-and MCL1-related metrics also showed appreciable correlations, the correlation values were lower, signaling that most AML cases examined relied on BCL2 for survival (Fig. 4b). Interestingly, the BCL2-BIM complex showed a much lower correlation than the BCL2-BAX complex count, reflecting our observation that the BCL2-BAX complexes’ dissociation predominantly mediated the pro-apoptotic effects of ABT-199 in HL60 cells (Fig. 4b-d). A simple combination of the four BCL2-related metrics only led to limited improvements in correlation while losing statistical significance, possibly due to redundant information concerning the BCL2 protein pool’s molecular status (Extended Data Fig. 8a,b and see below).

To explore metric combinations in a broader parametric space, we divided the 32 AML samples into responsive and non-responsive groups using an *ex vivo* AUC threshold of 0.61, corresponding to an IC_50_ of 100 nM ABT-199 (Extended Data Fig. 8c)^72,73^. We then examined approximately 10,000 different models, each with varying metric combinations and coefficients, selecting 67 models that made accurate predictions for the training set responsiveness, as validated by the remaining test samples (Extended Data Fig. 8d).

Remarkably, the final 67 models consistently identified a subgroup of selected metrics rather than displaying diverse metrics randomly (Fig. 4e). Two metrics, BCLxL-BIM_BH3_ PBA and BCLxL-BAK CPXs, were negatively correlated with *ex vivo* AUC values, presumably reflecting a compensatory role for BCLxL in counteracting accumulated apoptotic pressure in AMLs.

Upon detailed examination of the 67 models, we noticed a pattern where the information from each metric displayed minimal overlap. For example, while combining the BCL2-BAX CPX and BCL2-BIM_BH3_ PBA data successfully (which represented occupied and unoccupied portions of BCL2), neither metric could be combined with the BCL2 total level data (which included both occupied and unoccupied pools), likely due to partial overlap in the information they provided (Extended Data Fig. 8a). Similarly, the BCLxL-BAK CPX consistently exhibited higher absolute values in its correlations to *ex vivo* AUC than BCLxL-BAX CPX, possibly because of redundancy in the information concerning BAX that was already considered by the inclusion of the BCL2-BAX CPX metric (Fig.4e). As a result, a combination of BCL2-BIM_BH3_ PBA, BCL2-BAX CPX, and BCLxL-BAK CPX (with a negative coefficient) was selected, showing a slight but significant enhancement in the correlation to AUC values with statistical significance (*p*-value<0.05) for all metrics examined (Fig. 4f). In the receiver operating characteristic (ROC) analysis for binary prediction (responsive versus non-responsive), the resulting AUC showed the highest values for the combined model compared to single-metric models (Fig. 4g,h). Moreover, the mean squared error (MSE) was determined to be the smallest for the combined model, indicating higher accuracy in estimating ABT-199 efficacy (Fig. 4h).

We tested whether the created model could illustrate changes in the BCL2 PPI network as AML underwent clonal cancer evolution. For instance, BC-7064, a primary AML case, initially responded favorably to combined treatment with ABT-199 and azacitidine but relapsed after discontinuing the treatment (Fig. 4i). We performed PPI profiling on the initial (BC-7064) and relapsed (BC-7064-R) samples using our platform in a retrospective manner, comparing the three diagnostic PPI metrics in the combined model (Fig. 4i). The initial sample exhibited parameters indicative of strong ABT-199 effectiveness (estimated score of 0.78), composed of upregulated BCL2-BIM_BH3_ PBA and BCL2-BAX CPX counts and a low BCLxL-BAK CPX level (Fig. 4j). Conversely, the two BCL2-related metrics were more than halved for BC-7064-R, signaling a reduced driving force for ABT-199 efficacy. Although the resistance factor BCLxL-BAK complex also decreased for BC-7064-R, the weakened BCL2 dependence led to an estimated score of 0.36, explaining the relapse of this specific blood cancer (Fig. 4i,j).

Finally, we sought to make quantitative comparisons between our BCL2 PPI profiling and other established methods for predicting BH3-mimetic effectiveness, such as BH3 profiling^74,75^. BH3 profiling involves the addition of BH3 peptides derived from BIM, BAD, and HRK to individual AML samples after cell permeabilization, with mitochondrial depolarization monitored using JC-1 dye fluorescence (Extended Data Fig. 9a, upper). In our study, HL60 cells demonstrated robust depolarization signals in response to the BAD peptide, but not to the HRK peptide, a finding consistent with prior research (Extended Data Fig. 9b,c)^52,76^. We continued by examining 14 primary AML samples from the same cohort (due to limited sample availability) and tested various combinations of the depolarization signals elicited by BIM, BAD, and HRK peptides at different concentrations (Supplementary Table 3). We found that a combination that examined the difference between BAD and HRK signals (both measured at 10 μM peptides) yielded the highest correlation value of 0.65 with *ex vivo* ABT-199 efficacy (Extended Data Fig. 9d). Importantly, for this smaller cohort consisting of 14 AML samples, the BCL2 PPI profiling essentially maintained the performance parameters examined, including correlation, MSE, and AUC from the ROC analysis, which were superior to those determined for the BH3 profiling (Extended Data Fig. 9d-g). Attempts to use the BCL2 level measured by flow cytometry as a predictive marker (Extended Data Fig. 9a, lower) only resulted in muted performance across all evaluated parameters, further emphasizing the effectiveness of our profiling technique in estimating ABT-199 efficacy (Extended Data Fig. 9h-j).

### Construction of an MCL1 inhibitor efficacy model

Next, we investigated the possibility of training an analysis model for the MCL1 inhibitor AZD-5991, employing a similar process to that which we established for ABT-199. For this purpose, we evaluated the *ex vivo* efficacy of AZD-5991 on the same primary AML cohort (27 out of 32 total samples), comparing the determined AUC values with the BCL2 family metrics outlined in Figure 4 (Supplementary Table 2). Both the total MCL1 level and the MCL1 PBA count displayed correlation with AZD-5991 efficacy, though the correlations were diminished (Fig. 5a-c). Notably, the BIM_BH3_ PBA assay for BCLxL exhibited a negative correlation with AZD-5991 efficacy (Fig. 5d). Plotting the BCLxL PBA counts for responsive (*ex vivo* AUC≥0.61) versus non-responsive samples concerning AZD-5991 revealed that the non-responsive group manifested distinctly higher counts^72,73^, indicative of upregulated activities of BCLxL proteins within this group (Fig. 5e and Extended Data Fig. 8f). These collective findings suggest that BCLxL proteins play a substantial compensatory role when MCL1 proteins are saturated with inhibitor molecules. Furthermore, MCL1 protein complexes presented only limited correlations to *ex vivo* AUC, aligning with our hypothesis presented in Figure 3 that AZD-5991 may operate by preventing additional binding of pro-apoptotic proteins to MCL1 (i.e., saturation of MCL1 proteins), rather than inducing dissociation of pre-existing MCL1 protein complexes (Fig. 5a).

**Fig. 5.**
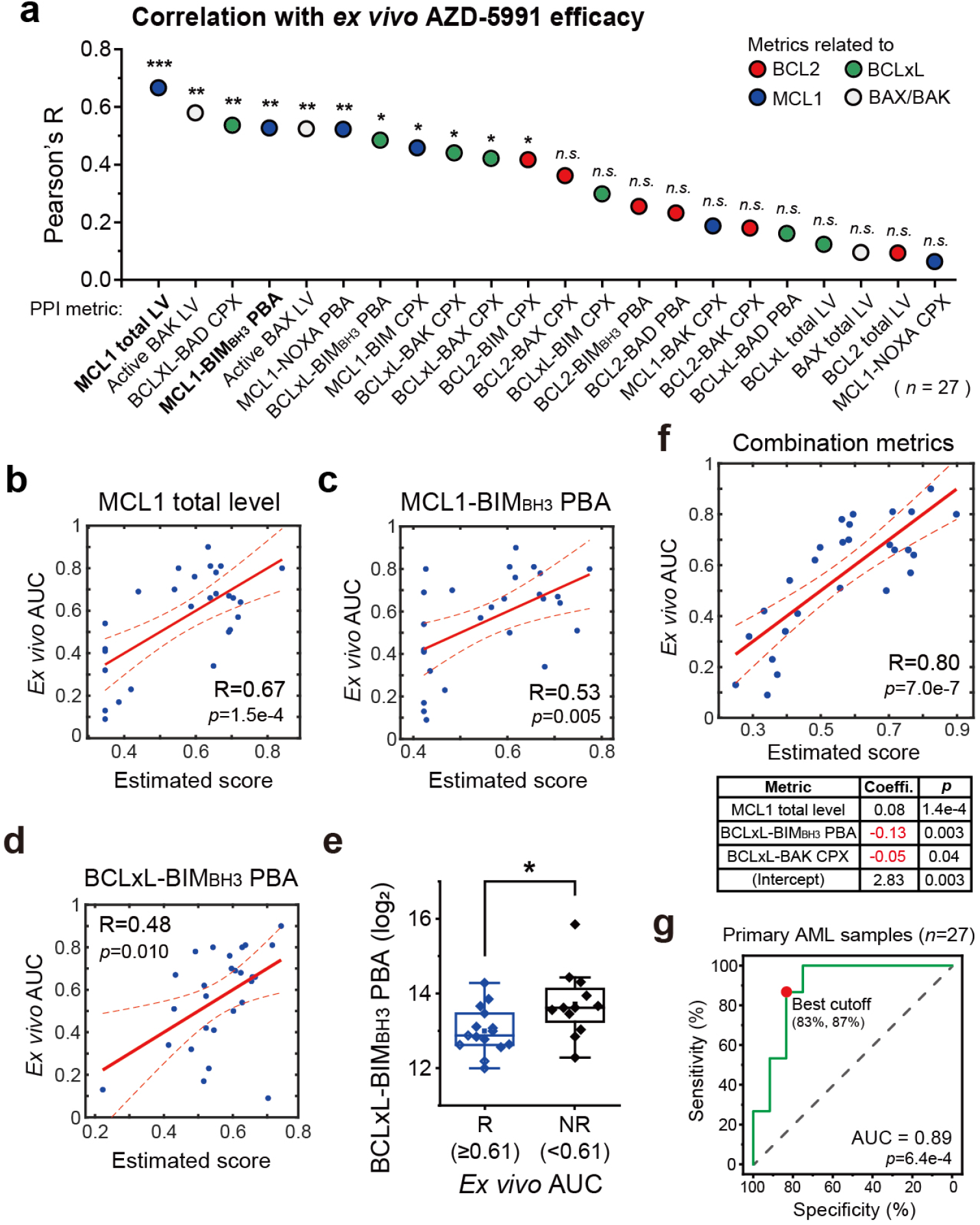
| Construction of an MCL1 inhibitor efficacy model. **a**, Pearson correlations between *ex vivo* AUC of AZD-5991 and PPI profiles for primary AML samples (*p-value*: *<0.05, **<0.01, ***<0.001, *n.s.*=not significant, *n*=27). **b-d**, Correlations between *ex vivo* AUC and single PPI metrics. (**b**) MCL1 total level (coefficient: 0.05), (**c**) MCL1-BIM_BH3_ PBA (coefficient: 0.04), (**d**) BCLxL-BIM_BH3_ PBA (coefficient: -0.13). **e**, Comparison of BCLxL-BIM_BH3_ PBA counts for the AZD-5991 to responsive (*ex vivo* AUC≥0.61, *n*=12) and non-responsive (*ex vivo* AUC<0.61, *n* =15) AML samples (Two-sided two-sample *t*-test, *p*-value=0.03). **f**, Correlation between *ex vivo* AUC and combination of multiple PPI metrics (MCL1 total level, BCLxL-BIM_BH3_ PBA, BCLxL-BAK CPX). The statistical indicators (coefficient and *p*-value) of each metric were presented. **g,** ROC curve between the estimated score and the *ex vivo* efficacy.

Integrating multiple parameters, namely the MCL1 total level, BCLxL-BIM_BH3_ PBA, and BCLxL-BAK CPX, we formulated a linear regression model that yielded an optimized correlation with the *ex vivo* AZD-5991 efficacy (Fig. 5f). While several BCL2-related parameters alone already displayed high correlations with ABT-199 efficacy (signaling the singular dependence on BCL2) (Fig. 4b), the MCL1-related metrics required combination with BCLxL-involving metrics to enhance the correlation, pointing to a complex interplay and dynamic between MCL1 and BCLxL protein pools (Fig. 5a,f and Extended Data Fig. 8e). The principle of minimal information overlap between metrics appeared to be preserved here as well, given that the final combination encompassed occupied and unoccupied populations of both MCL1 and BCLxL with seemingly minimal overlap. Through the combined model for AZD-5991 effectiveness, we achieved a high accuracy amounting to an AUC of 0.89 and statistically significant discriminative performance for binary prediction with ROC analysis (Fig. 5g). Altogether, our multi-parameter PPI data not only facilitates the construction of an analysis model for the efficacy of BH3 mimetics, but also allows us to identify the relative contributions of each assay data and the intricate balances among the protein complexes, potentially shedding light on the drugs’ modes of action.

### Using the PPI analysis model as a predictive biomarker for ABT-199 efficacy

We next explored whether BCL2 PPI profiling could proactively guide therapeutic decisions. BCL2 PPI profiling was performed for ten patients who were treated with ABT-199 through oral administration. Treatment outcomes were evaluated in accordance with European LeukemiaNet (ELN) recommendations^77^, and sequential samples were collected throughout the treatment journey (Fig. 6a and Supplementary Table 4).

**Fig. 6.**
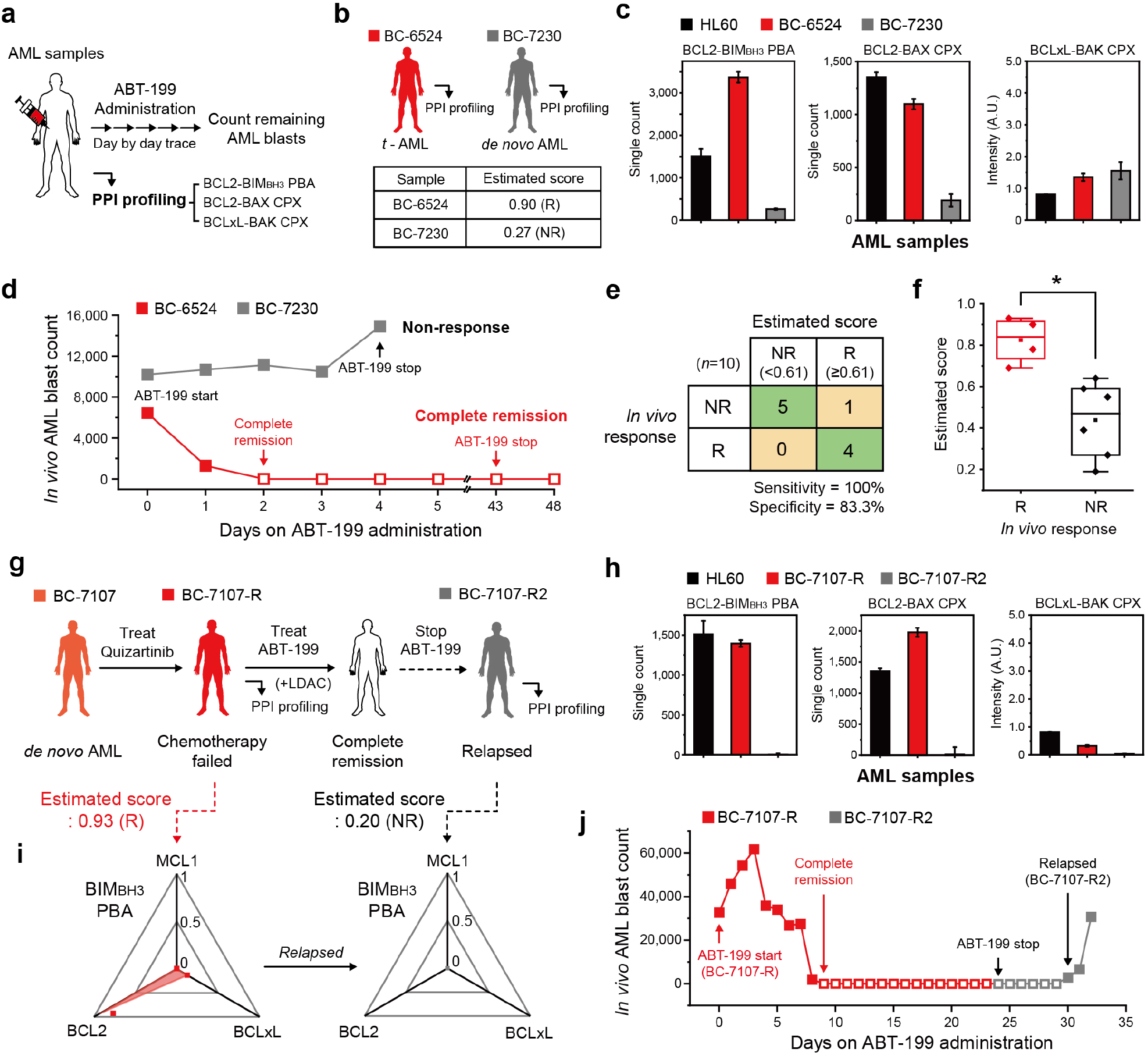
| Using the PPI analysis model as a predictive biomarker for *in vivo* ABT-199 response. **a**, Schematics of the study. Complete blood cell count (CBC) analysis was carried out on a daily basis to count the number of AML blasts in the primary AML samples. **b-d**, Comparison of the PPI profiles from the BC-6524 and BC-7230 samples. (**b**) Clinical features and the estimated scores with PPI diagnostic results (R: responsive, NR: non-responsive) for ABT-199, (**c**) Comparison of BCL2 family PPI profiles, (**d**) Changes of *in vivo* AML blast counts through the days after ABT-199 administration. **e**, Confusion matrix for the drug response prediction based on the estimated score and the *in vivo* drug responses (*n*=10). **f,** Comparison of the estimated scores of AML patients between responsive (*n*=4) and non-responsive (*n* =6) patients for *in vivo* ABT-199 administration (Two-sided Mann-Whitney test, *p*-value=0.014). **g**, Clinical features and the ABT-199 administration history of BC-7107. **h**, Tracking the changes of BCL2 family PPI profiles after relapse of BC-7107-R. **i**, Tracking the changes of BIM_BH3_ PBA profiles and PPI diagnostic results after relapse of BC-7107-R. **j**, Changes of *in vivo* AML blast counts from BC-7107-R through the days after ABT-199 administration. (**b**,**g**) The estimated scores were calculated from the model in Figure 4f. Error bars represent means ± s.d.

For example, two patients, BC-6524 and BC-7230, exhibited contrasting patterns in our PPI profiling, mirroring their disparate responses to ABT-199 treatment (Fig. 6b-d). Within the combination analysis model detailed in Figure 4f, BC-6524 was anticipated to respond to a BCL2-targeted inhibitor with a high estimated score of 0.90, while BC-7230 was predicted as a non-responder with an estimated score of 0.27 (Fig. 6b).

Specifically, BC-6524 displayed significantly higher counts for the BCL2-BIM_BH3_ PBA and BCL2-BAX CPX assays compared to BC-7230, while the BCLxL-BAK complex counts were low for both patients (Fig. 6c). Impressively, these diagnostic predictions were strongly corroborated by the *in vivo* responses. For patient BC-6524, peripheral blast counts began to decrease from day 2, culminating in confirmed complete remission as per bone marrow examination (Fig. 6d, red). Conversely, patient BC-7230 did not respond to ABT-199, as evidenced by increasing peripheral AML blasts from day 3 (Fig. 6d, grey).

When the 10 cases grouped into responsive (including complete remission and partial remission) and non-responsive categories, our predictions for responder (R) and non-responder (NR) with a threshold score of 0.61 aligned remarkably with the *in vivo* clinical outcomes for the nine patients (out of total ten patients), including those for BC-6524 and BC-7230 previously described, achieving 100% sensitivity and 83% specificity (Fig. 6d,e and Extended Data Fig. 10a-g). Moreover, the estimated scores for the initial responses between the two groups manifested a significant difference as determined by the Mann-Whitney test, which further increased when we included six additional samples from the same cohort collected after relapse (Fig. 6f and Extended Data Fig. 10h). An intriguing case was BC-7052, where our model initially projected a favorable score of 0.64 (Extended Data Fig. 10a). After a short initial response, the patient entered partial remission and ceased ABT-199 treatment. Interestingly, in *ex vivo* efficacy determination, the sample recorded a high AUC value of 0.63, precisely mirroring the prediction by the analysis model (Supplementary Table 4). This observation highlights a current model limitation, which was trained on *ex vivo* drug efficacy, and suggests that incorporating factors unique to *in vivo* cancer evolution may refine the model’s predictive capabilities. For example, the BCLxL PBA count, absent from the current model, was conspicuously high for BC-7052, hinting at a scenario where BCLxL proteins might become upregulated in activity as ABT-199 treatment persisted (Extended Data Fig. 10a, right).

In addition to the analysis of the BC-7064 case presented in Figure 4i, we conducted a longitudinal tracking study on another case, BC-7107, to gain insights into AML evolution during ABT-199 treatment (Fig. 6g). This patient, who had relapsed refractory disease and previously undergone treatment with another chemotherapeutic agent (Quizartinib), initially exhibited a robust BCL2 dependence (with an estimated score of 0.93) (Fig. 6g-i). Notably, the patient achieved complete remission upon combined treatment with ABT-199 and LDAC (Low-dose cytarabine) (Fig. 6g,j). However, after one month of treatment, relapse occurred (Fig. 6j). Interestingly, PPI profiling after the relapse showed nearly background signals for all three examined anti-apoptotic proteins, suggesting that the cancer might have developed a novel pathway to evade apoptosis initiation (Fig. 6h-j).

## Discussion

In our pursuit of establishing a precision medicine platform for protein complex-targeting therapies, we harnessed the capabilities of the SMPC platform to comprehensively profile BCL2 family protein complexes within specific AMLs. This endeavor involved optimizing extraction protocols applicable to clinical samples, ensuring the preservation of most protein complexes and retaining specific BCL2 family protein conformations through gentle lysis techniques. The integration of an anti-fouling polymer layer minimized cross-interactions and non-specific adsorption during surface pull-down procedures. Additionally, the direct single-molecule counting of target protein complexes demonstrated sustained sensitivity, effectively working even when only a few hundred protein complexes were immobilized per imaging area (∼10^4^ μm²). These technical innovations collectively enabled the assessment of over 20 distinct PPI species within clinical samples comprising as few as ∼1.2×10^6^ cells, achieving a level of sensitivity and accuracy amenable to clinical applications.

Given that BH3 mimetics function as PPI inhibitors, conventional genomic and proteomic profiling inherently falls short in capturing the dynamic rewiring of the PPI network and, consequently, predicting the response to specific BH3 mimetics^46–49,78,79^. Indeed, our findings demonstrated that the counts of BCL2 family protein complexes underwent drastic changes of more than three-to five-fold, while corresponding protein levels exhibited marginal alterations approximately less than 30%. Moreover, the effect of the two examined BH3 mimetics, ABT-199 and AZD-5991, on BCL2 family protein complexes appeared distinct, likely attributed to their distinct mechanisms of action (as discussed below). This underscores how relying solely on genomic mutations and proteomic levels, without considering their interconnectedness, may fail to fully elucidate the intricate complexity of BCL2 family biology underlying BH3 mimetic efficacy.

Presently, there are competing hypotheses regarding the primary species of protein complexes that BH3 mimetics target to dissociate for initiating apoptosis^16,80,81^. The activated BAX and BAK proteins, responsible for forming BAX-BAK pores and mediating MOMP, can either be newly expressed by cells or released from anti-apoptotic proteins. In the former scenario, BH3 mimetics saturate the target anti-apoptotic proteins, preventing further sequestration of nascent BAX and BAK proteins. In the latter, BH3 mimetics bind to the binding groove of anti-apoptotic proteins, causing the release of previously bound pro-apoptotic proteins. A detailed analysis of changes in relevant protein complexes and protein levels, along with their intricate balances, revealed that ABT-199 primarily acted by disassembling pre-existing BCL2 protein complexes. Conversely, the MCL1 inhibitor AZD-5591 appeared to exert its efficacy by interfering with the binding of newly produced BAK proteins to MCL1. Furthermore, tracing the behavior of the two types of BCL2 protein complexes over time indicated that the dissociation of BCL2-BAX complexes – not BCL2-BIM complexes – closely correlated with the onset of apoptosis in AML cells induced by ABT-199. Our findings are consistent with previous studies that employed fluorescence lifetime imaging microscopy to observe the release of BAD and BID, but not of BIM, from BCL2 and BCLxL by ABT-263^82^. Additionally, these studies noted pro-apoptotic effects triggered by the disruption of BCLxL-BAX complexes through BAD in isolated mouse liver mitochondria^83^. Indeed, in our multi-parametric measurement and linear regression analysis, the quantity of BCL2-BAX complexes was more closely tied to the *ex vivo* efficacy of ABT-199 than BCL2-BIM complexes. This led us to conclude that, in the specific AML cases studied, the dissociation of BCL2-BAX complexes primarily drove the efficacy of ABT-199.

These data compelled us to hypothesize a particular group of driver protein complexes that powerfully and positively correlated with the efficacy of the BH3 mimetics. We also found protein or PPI metrics associated with the drug efficacy with negative coefficients, which we defined as resistance complexes. The resulting analysis models included both driver and resistance complexes, reflecting the competing roles of different anti-apoptotic proteins in achieving the BH3-mimetic efficacy. In the combined model for ABT-199 effectiveness, BCL2-BIM PBA and BCL2-BAX CPX worked as the driver complexes, while BCLxL-BAK CPX worked as the resistance complex. However, the coefficient for BCLxL-BAK CPX was much smaller than those of BCL2-related parameters, consistent with the singular dependence of AMLs on BCL2. On the other hand, the analysis model for AZD-5591 efficacy, consisting of MCL1 total level, BCLxL-BAK CPX, and BCLxL-BIM_BH3_ PBA, assigned larger coefficients to two BCLxL-involved parameters than it did for the MCL1 total level, suggesting that MCL1 was in direct competition with BCLxL in mediating the effectiveness of AZD-5591. It is important to note that our current suite of assays lacks the capability to detect certain anti- and pro-apoptotic proteins in the BCL2 family, including BFL1 and BID. By extending our assay to incorporate these proteins – achievable through the screening of appropriate antibodies for IP and detection – we anticipate a more comprehensive profiling of the BCL2 family PPI network, as well as enhanced accuracy in our analysis model.

Our analysis model bears an intrinsic limitation as it was constructed based on drug efficacy determined *ex vivo*. Additionally, due to the relatively small size of the patient cohort employed in this study, the applicability of the model to larger cohorts remains untested. Notwithstanding these considerations, we took the step of applying the analysis model for prospective stratification of ABT-199 responders.

Remarkably, in nine out of ten cases, predictions based on three PPI metrics showed a significant correlation with clinical responses, achieving 100% sensitivity and 83.3% specificity. Additionally, our longitudinal tracking approach allowed us to observe how PPI connectivity evolved among the BCL2 family proteins, ultimately contributing to the development of resistance and cancer recurrence. While obtaining frozen BMMC or PBMC samples from hematological malignancies does not present significant challenges, we expect that the extraction of protein complexes from fixed tissues will be a crucial technical factor for successfully extending this method to solid tumors. Furthermore, ready access to the surface-passivated reaction chip and the single-molecule imaging device used in this study is essential for the widespread adoption of this diagnostic method in clinical decision-making.

In summary, our findings advocate for the SMPC platform as an effective method of scrutinizing multiple protein complexes in clinical samples with heightened sensitivity and precision. While emerging technologies such as single protein sequencing primarily serve as discovery tools^84^, the platform showcased in this study is tailored to the examination of dozens of protein complexes in clinical specimens. These profiles distill into information that is most directly pertinent to drug efficacies, thereby facilitating treatment decisions that would be considerably challenging to achieve through genomic and proteomic profiling alone. Given the rise in protein complex-targeting therapies, it will be intriguing to see whether the methodologies established herein could be extended to address other disease-associated protein complexes and drugs beyond the BCL2 family and BH3 mimetics.

## Methods

### Antibodies

Anti-rabbit immunoglobulin G (IgG) with biotin conjugation (111-065-144; Jackson ImmunoResearch) and anti-mouse immunoglobulin G (IgG) with biotin conjugation (715-066-151; Jackson ImmunoResearch) were used to immobilize the antibodies for surface IP. Anti-RFP antibody with biotin conjugation (ab34771; Abcam) and anti-GFP antibody with biotin conjugation (ab6658; Abcam) were used to immobilize mCherry-or eGFP-labeled proteins on surface. Anti-MCL1 (94296S; Cell Signaling Technology), BCLxL (MA5-15142; Thermo Fisher Scientific), BCL2 (4223S; Cell Signaling Technology), BAX (5023S; Cell Signaling Technology), and BAK (ab32371; Abcam) antibodies were used to immobilize the corresponding proteins via surface pull-down. Anti-MCL1 (MAB8825; Abnova), BCLxL (NBP1-47665; Novus Biologicals), BCL2 (sc-7382; Santa Cruz Biotechnology), BAX (MABC1176M; Sigma Aldrich), BAK (sc-517390; Santa Cruz Biotechnology), BIM (sc-374358; Santa Cruz Biotechnology), BAD (sc-8044; Santa Cruz Biotechnology), NOXA (sc-56169; Santa Cruz Biotechnology) antibodies were used to detect the corresponding proteins in the protein total level and the protein complex assays. To measure BCL2-BAX complex level, anti-BAX (5023S; Cell Signaling Technology) and anti-BCL2 (BMS1028; Invitrogen) antibodies were used to immobilize and detect the BCL2-BAX complex, respectively. Anti-rabbit immunoglobulin G (IgG) with Cy3 conjugation (111-165-046; Jackson ImmunoResearch) and anti-mouse IgG with Cy3 conjugation (715-165-151; Jackson ImmunoResearch) were used to label the detection antibodies for immunoassay. Anti-BCL2 antibody (BMS1028; Invitrogen), mouse IgG1 kappa isotype control (554121; BD Biosciences), and anti-mouse immunoglobulin G (IgG) with PE conjugation (715-116-150; Jackson ImmunoResearch) were used to detect the protein levels for quantitative flow cytometry.

### Drug reagents

Staurosporine (HY-15141; MedChemExpress), ABT-199 (HY-15531; MedChemExpress), and AZD-5991 (C-1060; Chemgood) were used to treat the blood cancer cell lines or primary AML cells. The drugs were diluted to 10 mM with DMSO and stored at -80 °C.

### Cell culture

HL60, THP-1, U937, and Ramos cells were purchased from Korean Cell Line Bank. HEK293T and SU-DHL8 cells were purchased from American Type Culture Collection (ATCC). NB4 cells were purchased from German Collection of Microorganisms and Cell Cultures (DSMZ). PC9 cells were provided by Y. Hayata (Kyushu University Faculty of Medicine, Japan). HEK293T cells were cultured in DMEM (D6429; Sigma Aldrich) supplemented with 10% (v/v) FBS (26140-079; Gibco) and 100 μg/ml penicillin/streptomycin (15140-122; Gibco). HL60, THP-1, NB4, U937, Ramos, SU-DHL8, and PC9 cells were cultured in RPMI1640 (R8758; Sigma Aldrich) supplemented with 10 % (v/v) FBS and 100 μg/ml penicillin/streptomycin. For THP-1, 0.05 mM of 2-mercaptoethanol (M3148; Sigma Aldrich) was added to culture media. All cells were incubated in a humidified incubator at 37 °C, 5% CO_2_. HEK293T and PC9 cells were rinsed with cold DPBS (D8537; Sigma Aldrich) before collection. Cells were collected with the scraper (90020; SPL Life Sciences) in 1 ml of cold DPBS. Suspension-type cells were collected with centrifugation at 500 g for 5 minutes at 4 °C and rinsed with 1 ml of cold DPBS. The cell suspensions were centrifuged at 500 g for 5 minutes at 4 °C. The supernatants were discarded after centrifugation and the cell pellets were stored at -80 °C by snap-freezing with liquid nitrogen.

### Primary AML samples

The cryo-preserved bone marrow mononuclear cells (BMMCs) and peripheral blood mononuclear cells (PBMCs) collected from AML patients treated at Seoul National University Hospital were used. Samples were collected between January 2014 and December 2019. Acute promyelocytic leukemias and biphenotypic leukemias were excluded. Mononuclear cells were isolated by ficoll gradient centrifugation. All samples were frozen in liquid nitrogen before flow cytometry and SMPC analysis. Aforementioned cell line collection protocol was applied for primary AML cell collection. Culturing method for flow cytometry assay is described on the sub-section on Drug treatment and drug efficacy measurement. Detailed information of each sample is described in Supplementary Table 1. This study was conducted according to the Declaration of Helsinki and was approved by the institutional review board of Seoul National University Hospital (IRB number: H-1910-176-107).

### Fluorescence-labeled protein constructions

All full-length BCL2 family proteins (*BCL2* (HG10195-M; SinoBiological), *BCLxL* (HG10455-M; SinoBiological), *MCL1* (HG10240-M; SinoBiological), *BIM_EL_* (HG13816-G; SinoBiological), *BAD* (HG10020-M; SinoBiological), *BAX* (HG11619-M; SinoBiological), *BAK* (HG10450-M; SinoBiological) and *NOXA* (HG16548-U; SinoBiological)) were isolated from their respective human cDNA. cDNA of *BIM_BH3_* (MRQAEPADMRPEIWIAQELRRIGDEFNAYYARR) was isolated from *BIM_EL_* cDNA. cDNA of *BAK fragments* – residues 24-69 (α_1_ helix), 70-91 (α_2_-α_3_ helices), 90-150 (α_3_-α_5_ helices), and 146-187 (α_6_-α_8_ helices) – were isolated from *BAK* cDNA. All cDNAs were cloned into *pCMV* vectors with either eGFP or mCherry sequences to generate fluorescence-labeled proteins using Gibson assembly. The fluorescence proteins were fused to the carboxyl-end of BCL2 family proteins. The resulting plasmids were introduced into HEK293T cells through transient transfection. Plasmid DNA with the amount of 10 μg was mixed with 30 μg of linear polyethyleneimine (PEI) (23966-100; Polysciences) in 1 ml of serum-free DMEM. The mixture was introduced into ∼2×10^6^ of HEK293T cells in a 90 mm^2^ cell culture plate.

### Cell lysis

All collected cell pellets were resuspended with the lysis buffer (0.2% or 1% (v/v) detergent, 50 mM HEPES with pH 7.4, 150 mM NaCl, 10% (v/v) glycerol, 1 mM EDTA, 2% (v/v) protease inhibitor cocktail (P8340; Sigma Aldrich), 2% (v/v) phosphatase inhibitor cocktail 2 (P5726; Sigma Aldrich), and 2% (v/v) phosphatase inhibitor cocktail 3 (P0044; Sigma Aldrich). Triton-X100 (X100; Sigma Aldrich), glyco-diosgenin (GDN) (GDN101; Anatrace), digitonin (D141; Sigma Aldrich), Tween 20 (P2287; Sigma Aldrich), 3-[(3-Cholamidopropyl) dimethylammonio]-1-propanesulfonate hydrate (CHAPS) (C3023; Sigma Aldrich), and sodium dodecyl sulfate (SDS) (L3771; Sigma Aldrich)) were used for optimizing the lysis protocol. Triton-X100, GDN, digitonin, Tween 20, and CHAPS were used in 1% (v/v), and SDS was used in 0.2% (v/v) in lysis buffer. HEK293T cells expressing probes for the PBA experiments – BIM_BH3_-eGFP, BIM_EL_-eGFP, BAD-eGFP, and NOXA-eGFP – were lysed with Triton-X100 lysis buffer. HEK293T cells expressing fluorescence protein labeled BCL2 family proteins (BCL2-mCherry, BCLxL-mCherry, MCL1-mCherry, BCL2-eGFP, BCLxL-eGFP, MCL1-eGFP, BAX-eGFP, BAK-eGFP, and BAK fragments-eGFP) were lysed with GDN lysis buffer for immunoassay and PBA. The construct designs and protein expressions in HEK293T cells for fluorescence protein labeled BCL2 family proteins are described on above. All the cancer cell lines, and primary AML cells were lysed with GDN lysis buffer for immunoassay and PBA.

For the cell lysis, the cell pellets were resuspended with the designated lysis buffer and incubated for 30 minutes at 4 °C. After lysis, the cell suspensions were centrifuged at 15,000 g for 10 minutes at 4 °C. The supernatants were isolated after centrifugation. The total protein concentration for each supernatant was measured with a DC protein assay kit (5000113, 5000114, 5000115; Bio-Rad) following the manufacturer’s instructions. The total concentration of the fluorescence proteins in each supernatant was measured with a Sense microplate reader (425-301; HIDEX). To quantify the fluorescence proteins, a laser wavelength of 485 nm was used for eGFP excitation and 544 nm was used for mCherry excitation respectively. Serial dilution experiments using eGFP and mCherry were performed to generate the calibration curves. The aforementioned calibration curves were used to quantify the concentrations of fluorescently labeled proteins. The supernatants were then aliquoted and stored at -80 °C followed by snap-freezing with liquid nitrogen.

### Drug treatment and drug efficacy measurement

All blood cancer cells and primary AML cells were precultured in a 25T flask (non-treated surface) for 3 hours before drug treatment. All cells were seeded in 4.5 ml of culture media and cultured in a humidified incubator at 37 °C, 5% CO_2_. STS, ABT-199, and AZD-5991 were then treated to cells at various concentrations. All drugs were initially diluted with DMSO to 1 mM. For experiments, serial dilution was performed with non-serum RPMI1640 to reach the target concentration. To determine the efficacy of drugs in blood cancer cells and primary AML cells, an AnnexinV/PI apoptosis assay was performed, where cells were stained with FITC-AnnexinV (640906; Biolegend) and PI (421301; Biolegend). After drug treatment, 5×10^5^ of cells were rinsed with cold DPBS and collected by centrifugation with 500 g for 5 minutes at 4 °C. After the supernatants were discarded, the cell pellets were resuspended with 100 μl of cold AnnexinV-binding buffer (422201; Biolegend). After resuspension, 10 μl of FITC-AnnexinV solution and 5 μl of PI solution were added into the cell suspensions. The cell suspensions were incubated for 15 minutes at room temperature avoiding exposure to light. After that, 400 μl of cold AnnexinV-binding buffer was added to avoid overstaining. The stained cells were analyzed with a flow cytometry (SH800S; Sony). The proportions of AnnexinV^-^/PI^-^ cells from each sample were calculated and converted to viability (%). Based on the measured viability, the drug efficacy curve was fitted using the *logistic* function inside the analysis software (OriginPro 2022), and the area under curve (AUC) was measured from the fitted curve.

### Single-molecule pull-down and co-IP for BCL2 family proteins

5 μg/ml of NeutrAvidin (31000; Thermo Fisher Scientific) was loaded into each reaction chamber of the Pi-Chip (PROTEINA) and incubated for 10 minutes. The imaging chip was washed with TritonX-100-buffer (0.1% (v/v) TritonX-100, 50 mM HEPES with pH 7.4, 150 mM NaCl) to remove unbound NeutrAvidin. Biotin-conjugated IgGs (anti-rabbit IgGs) were loaded to each reaction chamber with 1:200 in TritonX-100-buffer and incubated for 10 minutes, followed by washing with TritonX-100-buffer. Then, the previously mentioned monoclonal antibodies (anti-MCL1, BCLxL, BCL2, BAX, and BAK antibodies for surface IP) were introduced into each reaction chamber at a dilution of 1:100 in TritonX-100-buffer, and then incubated for a duration of 10 minutes. The imaging chip was washed with GDN-buffer (0.01% (w/v) GDN, 50 mM HEPES with pH 7.4, 150 mM NaCl, 1% (v/v) Glycerol, 1 mM EDTA). After washing, crude cell extracts were diluted based on the total protein concentrations and were loaded into each reaction chamber. During the process of immunoprecipitating fluorescence-labeled proteins, anti-RFP or anti-GFP antibodies were introduced at a dilution of 1:200 in TritonX-100-buffer. This mixture was then incubated for a period of 20 minutes, as an alternative to using biotin-conjugated IgGs.

To measure the total level of BCL2 family proteins, cell extracts with a concentration of 1 mg/ml were introduced into each reaction chamber. These chambers were pre-coated with primary antibodies for immunoprecipitation, and the samples were allowed to incubate for 1 hour to capture of the target proteins. To measure the total level of BAX and BAK, the cell extracts were mixed with the activation buffer (2% (v/v) Triton X-100, 50 mM HEPES (pH 7.4), 150 mM NaCl, and 1 mM EDTA) with 1:1 for 30 minutes before pull-down. After activation, the cell extracts were loaded into each reaction chamber and incubated for 1 hour. To measure the PPI complex levels, 2 mg/ml of cell extracts were loaded into each reaction chamber with 2 hours of incubation period to pull-down the target PPI complexes.

After the surface pull-down of target proteins or protein complexes, the previously mentioned primary antibodies were introduced for the purpose of detecting either total protein levels (anti-MCL1, BCLxL, BCL2, BAX, and BAK antibodies for immunoassay) or protein complexes (anti-BAX, BAK, BIM, BAD, and NOXA antibodies for immunoassay). These antibodies were loaded into each reaction chamber at a dilution of 1:100 in TritonX-100-buffer and then incubated for a duration of 1 hour. To label the detection antibodies, Cy3-conjugated IgGs (anti-mouse IgGs) were introduced into each reaction chamber at a dilution of 1:1,000 in TritonX-100-buffer. These chambers underwent an incubation period of 20 minutes.

To measure the PBA, 1 mg/ml of cell extracts were loaded into each reaction chamber and incubated for 1 hour. After that, the imaging chip was washed with GDN-buffer. The HEK293T cells expressing PBA probes were lysed with TritonX-100 lysis buffer. The crude cell extracts containing the PPI probes were diluted to achieve the target eGFP concentration. Subsequently, they were incubated for a duration of 10 minutes with the immunoprecipitated bait proteins.

### Single-molecule fluorescence imaging

Single-molecule fluorescence signal was assessed using the Pi-View system (PROTEINA), which allows for single-molecule fluorescence imaging at two different excitation lasers, 488 nm and 532 nm^55,57,65^. The emitted photons from a single fluorophore were collected by employing a sensitive sCMOS (scalable complementary metal-oxide semiconductor) camera. An autofocus system provides high throughput image acquisition across the whole imaging chip that contained 40 reaction chambers. To determine the single-molecule counts for each assay, ∼10 different locations were imaged in a single reaction chamber. Each image was acquired for 5 frames with 100 ms time resolution, and those frames were averaged to generate a single snapshot, which reduced random noise and improved singal-to-noise ratio. Protocols for image processing were detailed in Extended Data Fig. 2. The current imaging system identifies and counts up to 12,000 single-molecule fluorescence spots within a 100×100 µm² field of view, an upper limit for the identification of diffraction-limited spots. When the number of such spots exceeds this limit, overlapping begins to occur, compromising the ability to identify individual spots (Extended Data Fig. 2d). To address this limitation, the total fluorescence signals captured across all pixels within the field of view were integrated.

Simultaneously, by recording time-traces of photobleaching, a relationship between the number of single-molecule spots and the total fluorescence intensity was established, particularly, in the range that permitted the identification of individual spots (i.e., below the upper limit), which was typically linear (Extended Data Fig. 2e,f). This linear relationship ultimately enabled extrapolation of total fluorescence intensity into corresponding single-molecule counts in regions exceeding the upper limit (Extended Data Fig. 2g). The computational analysis codes are available on our GitHub.

### Drug efficacy prediction model analysis

Drug efficacy prediction models for AML patients were generated using linear regression analysis by MATLAB 2021a. The protein and protein complex data from the primary AML cells were all converted to a log_2_ scale. Any instance where the number of single-molecule fluorescence spots was less than 100 counts were fixed to the minimum value of 100 counts for log transformation. For BCLxL-BAX, BCLxL-BAK, and BCL2-BAK protein complexes, the total fluorescence signal data were used instead of single-molecule spot data, and the fluorescence signals less than 10^7^ were fixed to the minimum value (10^7^ A.U.) for log transformation. The *ex vivo* drug efficacy (*ex vivo* AUC) of the primary AML cells were obtained from the vaibility measuring method described on the sub-section on Drug treatment and drug efficacy measurement. The area under curves (AUCs) were initially calculated from the fitted curves for viability after BH3 mimetic treatment, and *ex vivo* drug efficacy were normalized by *ex vivo AUC=1-(AUC/Maximal AUC)*. The correlation between the PPI profiling data and the *ex vivo* drug efficacy was calculated through linear regression with the custom analysis code. Each generated model provided the statistical indicators, *p*-values for the linear regression itself, as well as for the intercept. The models were accepted only when their *p*-values were statistically significant (*p*-value<0.05). To construct combined metrics models, various randomly selected combinations of PPI profiles were assessed in order to identify a model that met two criteria: high correlation (indicated by a high R value) and statistical significance (with *p*-values < 0.05). Lasso regression analysis for selecting metrics which are highly correlated with drug response were generated by Python 3.11. The training and test groups were randomly selected from the primary AML sample cohort. The Lasso regression models were trained using the training group and evaluated based on Pearson’s R as well as prediction outcomes for the test group, and 67 models were selected from 10,000 different initial models (Extended Data Fig. 8d). The custom analysis codes and raw data utilized for generating the models can be accessed on our Github.

### BH3 profiling

We followed the methods outlined in the previous studies for BH3 profiling^74,75^. The MEB buffer (150 mM mannitol (M9647; Sigma Aldrich), 10 mM HEPES pH 7.4, 50 mM KCl (P9541; Sigma Aldrich), 0.02 mM EGTA (BE004; Biosolution), 0.02 mM EDTA, 0.1% (w/v) BSA (A4737; Sigma Aldrich), and 5 mM succinate (S3674; Sigma Aldrich)), along with the staining solution (20 μg/ml oligomycin (O4876; Sigma Aldrich), 50 μg/ml digitonin, 2 μM JC-1 (ENZ-52304; Enzo Life Sciences), and 10 mM 2-mercaptoethanol (M3148; Sigma Aldrich) in the MEB buffer) were prepared and stored at 4 °C. The BH3 peptides (or BH3 mimetics) were diluted with the staining solution to achieve twice the target concentration, producing the 2x staining solution for each peptide. The synthesized BH3 peptides (BIM (MRPEIWIAQELRRIGDEFNA, purity=98%), BAD (LWAAQRYGRELRRMSDEFEGSFKGL, purity=99%), and HRK (SSAAQLTAARLKALGDELHQY, purity=99%) all synthesized by PEPTRON (South Korea)), DMSO, carbonyl cyanide 4-(trifluoromethoxy) phenylhydrazone (FCCP) (C2920; Sigma Aldrich), ABT-199, and AZD-5991 were used for BH3 profiling. We aliquoted 50 μl of the 2x staining solution into a black-coated 96-well plate to produce a BH3 profiling plate, which was then stored at -80 °C until use.

To perform BH3 profiling for primary AML cells, 10^6^ of primary AML cells were rinsed with 1 ml of RPMI1640 supplemented with 10% (v/v) FBS. After centrifugation at 300 g for 5 minutes, the collected cells were resuspended in 1 ml of MEB buffer. Subsequently, the resuspended cells were then aliquoted into each well of the BH3 profiling plate that had been equilibrated to room temperature. The BH3 profiling plate was placed in a plate reader (Synergy H1; BioTek) set at a constant temperature of 32°C, and the fluorescence was measured every 5 minutes over a period of 3 hours. To measure the relative fluorescence units (RFU) at a wavelength of 590 nm, a light wavelength of 545 nm was used for JC-1 excitation. Based on the measured RFU, the area under the curve (AUC) was measured using the analysis software (OriginPro 2022). The depolarizations were calculated by the following equation, and the BH3 profiling results for primary AML cells can be found in Supplementary Information.

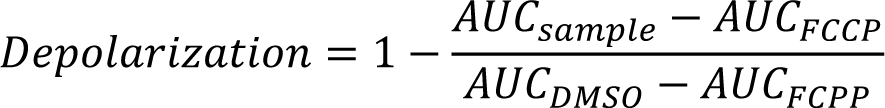

### Quantitative flow cytometry

We employed the methods outlined in previous studies to quantify protein levels using flow cytometry^85^. Briefly, 10^6^ of primary AML cells were rinsed with 1 ml of RPMI1640 supplemented with 10% (v/v) FBS and collected with centrifugation at 300 g for 5 minutes. The collected cells were resuspended in 0.5 ml of 37 °C lyse fix buffer (558049; BD Biosciences). The cell suspensions were centrifuged at 1,700 g for 5 minutes, after which the supernatants were discarded. The cell pellets were resuspended in staining wash buffer (0.5% (w/v) BSA and 0.1% (w/v) sodium azide (S2002; Sigma Aldrich) in DPBS) and aliquoted into two separate tubes at 100 μl each. 900 μl of permeabilization wash buffer (PW) (557885; BD Biosciences) was then added to each tube, followed by a 10-minute incubation at room temperature. Following the incubation, the tubes were subjected to centrifugation at 1,700 g for 5 minutes, and the supernatant was subsequently reduced to 100 μl. The permeabilized cells were stained with either 0.5 μg of anti-BCL2 antibody or an isotype control antibody respectively and incubated for 30 minutes. The cells were rinsed by adding 900 μl of PW and centrifuged at 1,700 g for 5 minutes. The rinsed cells were then stained with 1.5 μl of PE-conjugated anti-Mouse IgG and incubated for 30 minutes in a dark room to prevent light exposure. After rinsing to eliminate any unbound antibodies, the stained cells were resuspended in 350 μl of fix buffer (557870; BD Biosciences) for fixation and stored at 4°C until they were ready to be analyzed using a flow cytometer (SH800S; Sony). Quantum^TM^ R-PE MESF beads (827; Bangs Laboratories) were utilized to produce calibration curves. These curves were subsequently employed, following the manufacturer’s instructions, to quantify the levels of BCL2 in the samples.

### Reporting Summary

Further information on research design is available in the Nature Research Reporting Summary linked to this article.

### Data availability

The data supporting the findings of this study are available within the article and its Supplementary Information. All raw data generated or analysis during the study are available on the Github at the following link: https://github.com/tyyoonlab-snu/Nat-Biomed-Eng-2023-.

### Code availability

The custom MATLAB and Python codes used for drug efficacy prediction model analysis are available on the Github at the following link: https://github.com/tyyoonlab-snu/Nat-Biomed-Eng-2023-

## Acknowledgements

We thank Shi Ho Kim and Jason Sang Hyun Park for critical reading of the manuscript, and Changwon Kim, Gee Sung Eun, and Tae Gyun Kim for active discussion concerning data analysis. This work was supported by the National Grants for Leading Scientists (NRF-2021R1A3B1071354 to T.-Y.Y.) and the Bio Medical Technology Development Program (NRF-2018M3A9E2023523 to T.-Y.Y.) funded by the National Research Foundation of South Korea. This work was also supported by the Korea Health Technology R&D Project of the Korea Health Industry Development Institute (KHIDI) (HI14C1277 to Y.K.) funded by the Ministry of Health & Welfare of South Korea.

## Author contributions

T.-Y.Y. conceived of and supervised the project. C.C., J.M.B., M.C., H.L., B.C., Y.K., and T.-Y.Y. designed the experiments. C.C. and M.C. performed the single-molecule pull-down and co-IP experiments. C.C. and H.K. measured the inter-chip CVs of the immunoassays. J.M.B. and Y.K. collected and characterized primary AML samples and performed *in vivo* ABT-199 treatment journey. C.C., Y.L., and H.P. developed the PPI probes for PBA. C.C., M.C., B.C., H.K., S.H., and Y.L. performed the PPI profiling for the primary AML samples. C.C. and Y.L. performed BH3 profiling for the primary AML samples. C.C. and S.H. performed flow cytometry for the primary AML samples. C.C. and M.C. measured the *ex vivo* drug efficacies of the primary AML samples. H.L. developed the *ex vivo* drug efficacy predicting programs. C.C. analyzed and visualized all data. C.C. and T.-Y.Y. wrote the manuscript with inputs from all authors.

## Extended Data Figures

**Extended Data Fig. 1.**
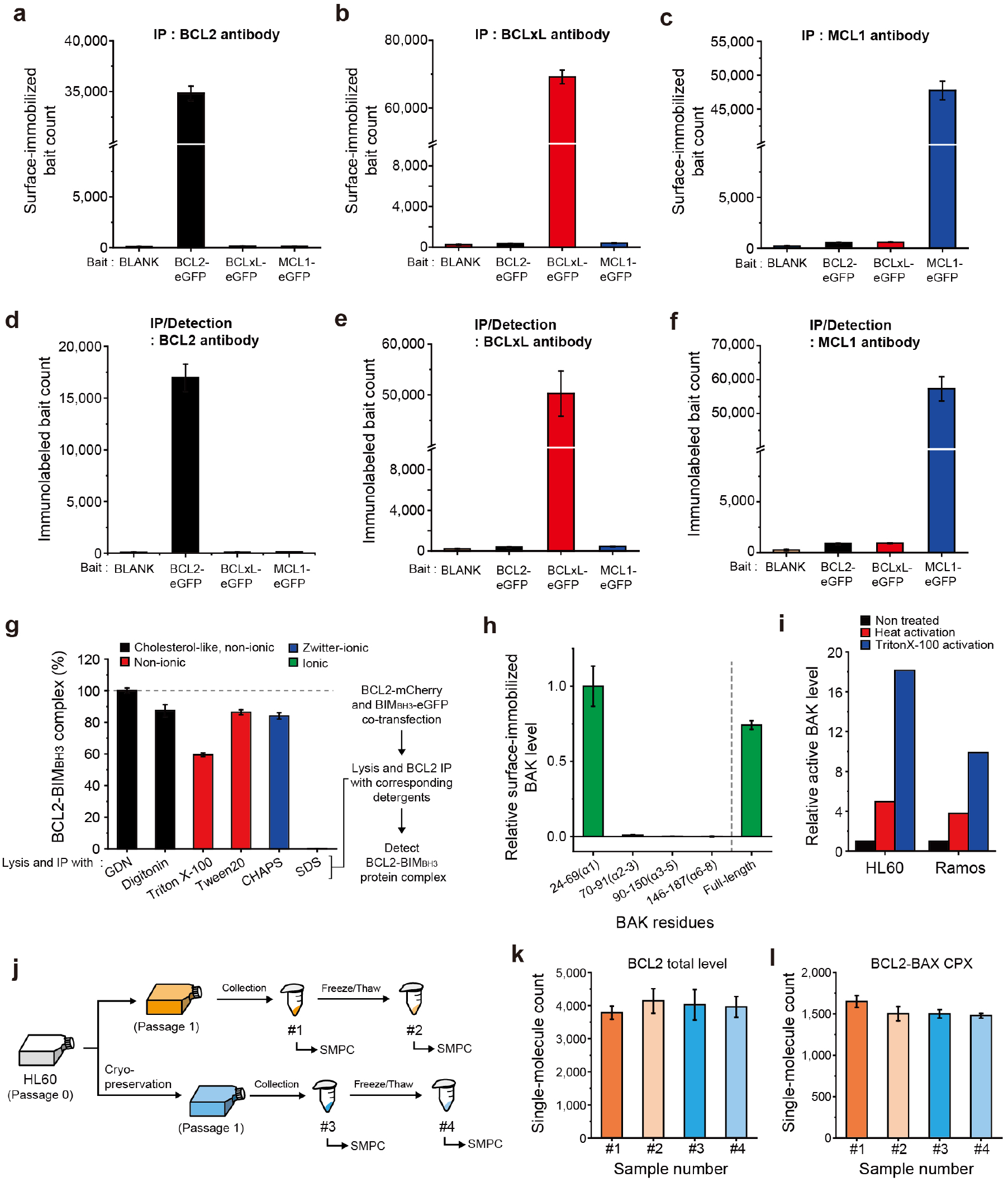
| Development of the single-molecule pull-down and co-IP (SMPC) platform to profile BCL2 family proteins. **a-c**, Selection of monoclonal antibodies for surface immunoprecipitation (IP). Surface IP of eGFP-labeled target anti-apoptotic proteins from transfected HEK293T crude cell extracts. Crude cell extracts with 1 nM of eGFP-labeled anti-apoptotic protein were used to IP. (**a**) BCL2 IP antibody, (**b**) BCLxL IP antibody, (**c**) MCL1 IP antibody. **d-f**, Selection of monoclonal antibodies for total level immunoassay of surface-immobilized target anti-apoptotic proteins. (**d**) BCL2 detection antibody, (**e**) BCLxL detection antibody, (**f**) MCL1 detection antibody. **g**, Relative model BCL2-BIM_BH3_ complex from BCL2-mCherry/BIM_BH3_-eGFP co-transfected HEK293T cells after the lysis with different detergent types. **h,** Detection of surface-immobilized BAK fragments by using the monoclonal antibody AT38E2. Crude cell extracts with 1 nM of eGFP-labeled BAK fragments were used to IP. **i,** Activation of BAK from HL60 and Ramos cells by heat or triton X-100 treatment to crude cell extracts^63,64^. **j,** Schematics for the preparation of HL60 samples with different preservation conditions. **k,l,** Comparison of the PPI profiles across the sample states using SMPC platform. (**k**) BCL total level, (**l**) BCL2-BAX complex. Error bars represent means±s.d.

**Extended Data Fig. 2.**
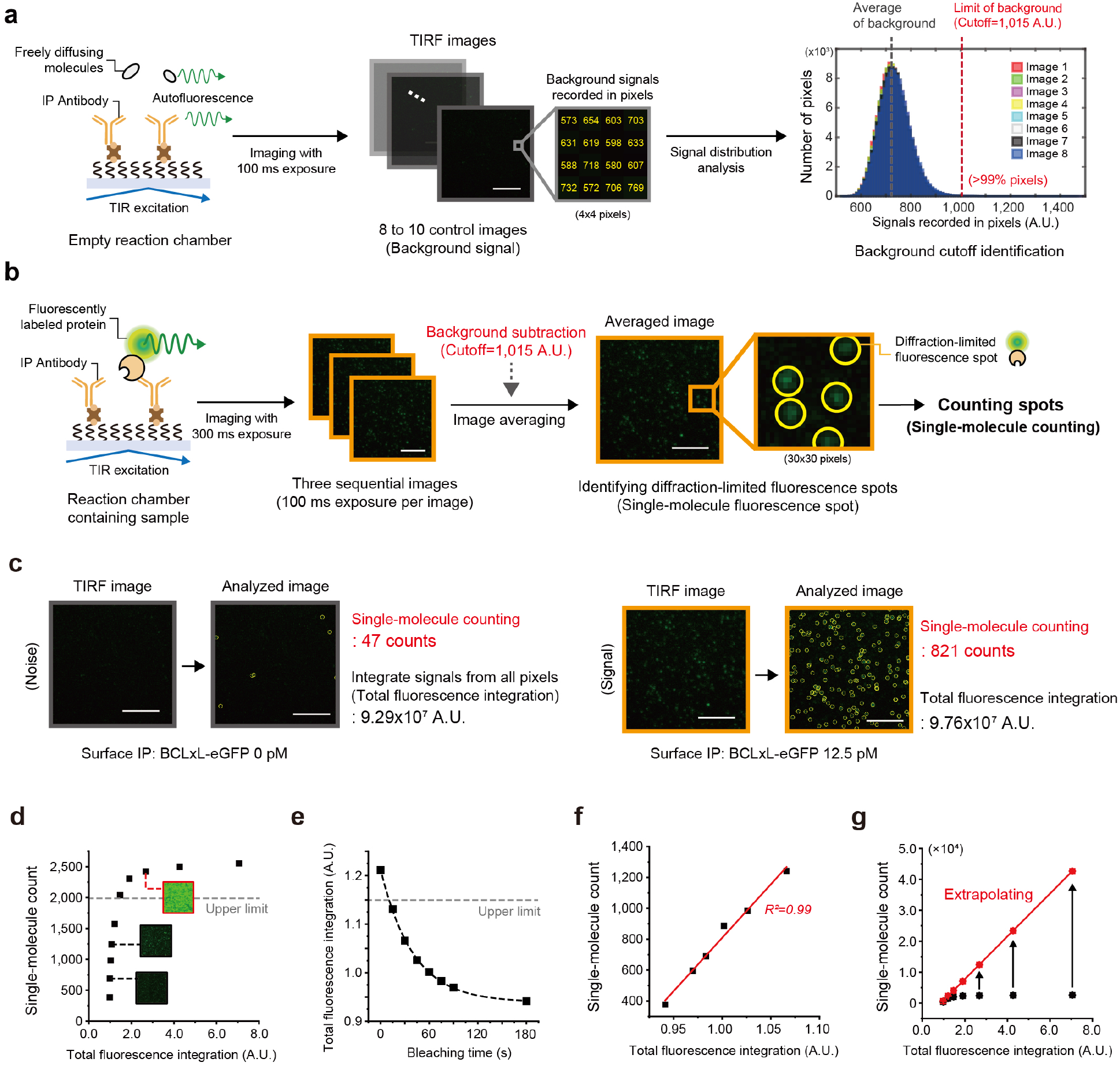
| Post image analysis for counting the number of single-molecule fluorescence spots. **a**, Setting the cutoff to eliminate background signals for post image analysis. We used 8 to 10 control images captured with a 100 ms time resolution from empty reaction chambers to establish a background limit. The background limit (cutoff=1,015 A.U.) was determined from the signal distribution recorded in each pixel of the control images, effectively eliminating 99% of background signals. **b,** Identification of diffraction-limited fluorescence spots. Three sequential images captured with a 300 ms time frame were averaged to create a single image upon subtracting the background. From the averaged image, the number of diffraction-limited fluorescence spots were counted (single-molecule counting). **c,** Identified single-molecule counts and integration of individual pixel signals of the imaging area (total fluorescence integration) from the analyzed TIRF images. (**a-d**) The TIRF images were obtained by surface IP of BCLxL-eGFP expressed in HEK293T cell extracts. (Scale bar: 10 μm). **d-g,** Calibration of single-molecule count from the total fluorescence integration. (**d**) Single-molecule count dependent on the total fluorescence integration of surface-immobilized BCL2-mCherry obtained from the analyzed images. (**e**) Photobleaching kinetics of surface-immobilized BCL2-mCherry signals depending on the bleaching time. (**f**) The linear correlation between single-molecule count and total fluorescence integration of surface-immobilized BCL2-mCherry after photobleaching. (**g**) Calibration of single-molecule count from total fluorescence integration by extrapolating. The linear correlation was obtained in (**f**). (**d-g**) The TIRF images were obtained by surface IP of BCL2-mCherry expressed in HEK293T cell extracts.

**Extended Data Fig. 3.**
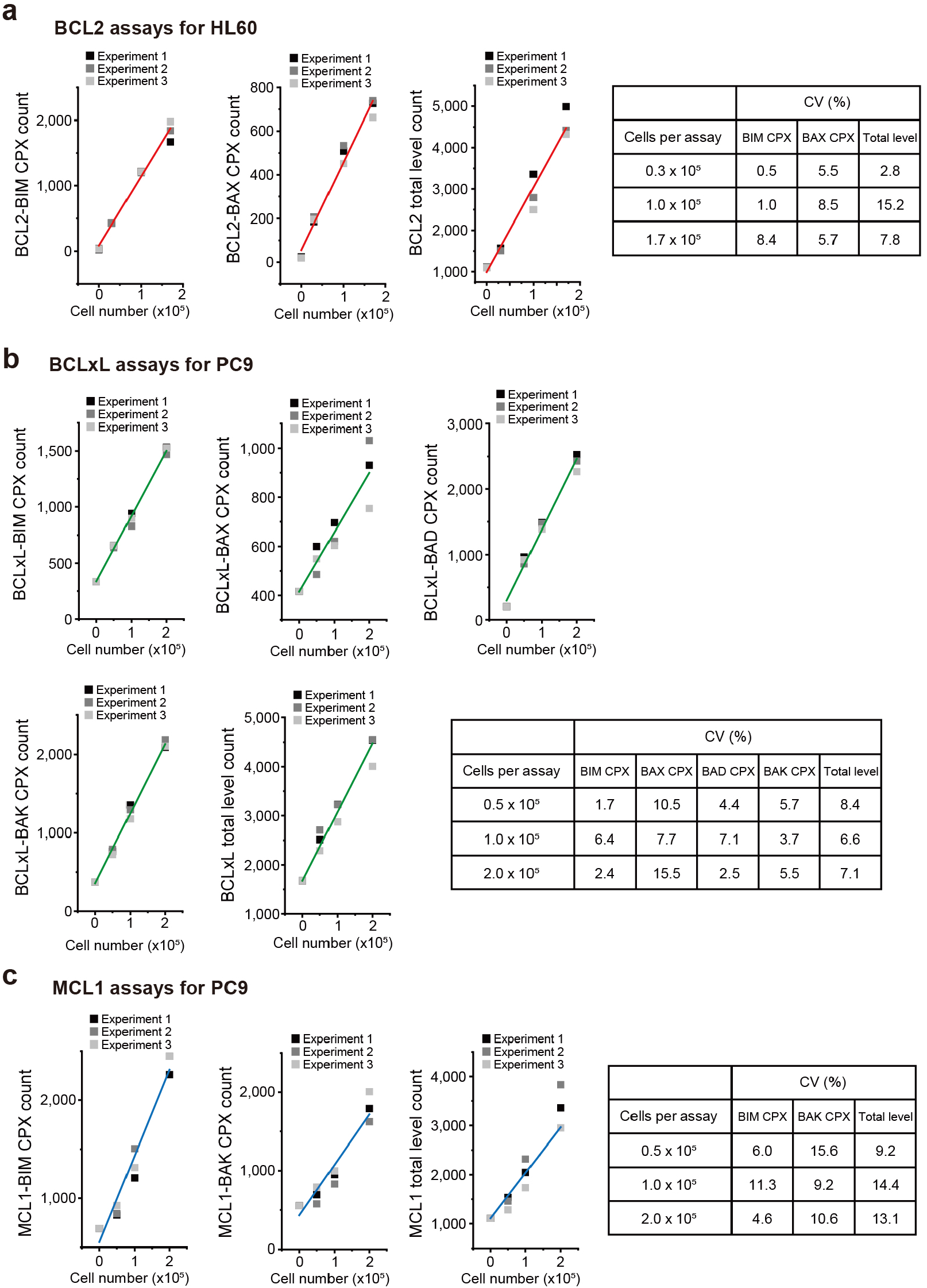
| Attesting the stability of the SMPC platform by inter-chip measurements of immunoassays. **a**, Counts of BCL2-related immunoassays (BCL2-BIM complex, BCL2-BAX complex, BCL2 total level) from the fixed numbers of HL60 cells (*n*=3). **b**, Counts of BCLxL-related immunoassays (BCLxL-BIM complex, BCLxL-BAX complex, BCLxL-BAD complex, BCLxL-BAK complex, BCLxL total level) from the fixed numbers of PC9 cells (*n*=3). **c**, Counts of MCL1-related immunoassays (MCL1-BIM complex, MCL1-BAK complex, MCL1 total level) from the fixed numbers of PC9 cells (*n*=3). The single-molecule counts were rescaled to account for the labeling efficiencies of the immunoassays as well as the specific incubation conditions for direct comparison. All data were measured from independent inter-chip experiments. CVs obtained from independent inter-chip measurement for all the immunoassays and cell numbers (*n*=3).

**Extended Data Fig. 4.**
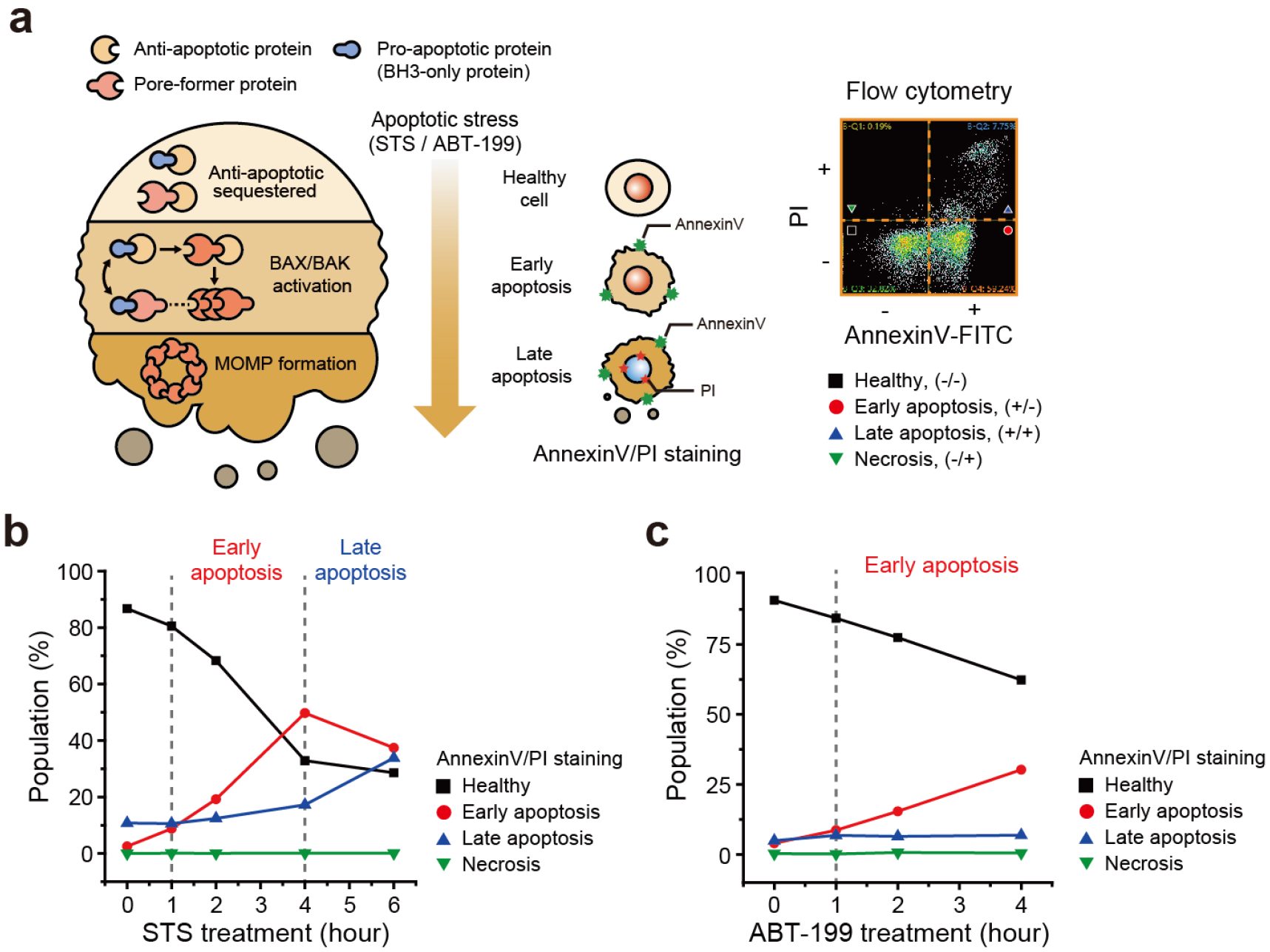
| Tracking the response of HL60 cells under apoptotic stress. **a**, Schematics for the intracellular BCL2-PPI profile changes and the result of flow cytometry analysis after Annexin V/PI staining through the progression of apoptosis pathway. **b**,**c**, Population changes of HL60 cells at different time points through the treatment of drug with Annexin V/PI staining. (**b**) 2 μM of STS, (**c**) 300 nM of ABT-199. Error bars represent means±s.d.

**Extended Data Fig. 5.**
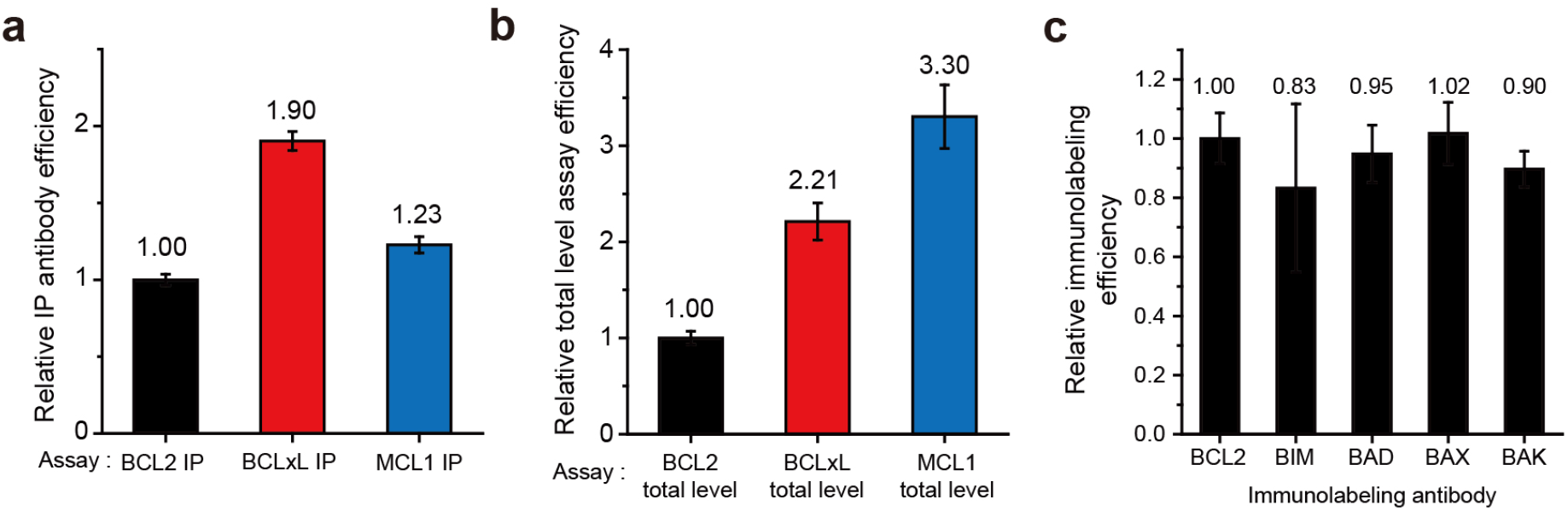
| Relatively comparing the normalized efficiencies of different immunoassays. **a**, Comparing relative efficiencies of IP antibodies for anti-apoptotic proteins. 1 nM of eGFP-labeled anti-apoptotic proteins were immobilized on the surface by the matched IP antibodies respectively, and the eGFP signals were compared. **b**, Comparing relative efficiencies of immunolabeling antibodies for anti-apoptotic proteins. The surface immobilized anti-apoptotic protein in (**a**) were immunolabeled by the matched labeling antibodies respectively, and the relative efficiencies for total level assays were compared. **c**, Comparing relative immunolabeling efficiencies of detection antibodies for BCL2 family proteins. 1 nM of eGFP-labeled BCL2 family proteins were immobilized on the surface by the anti-eGFP antibody and detected with matched detection antibodies for immunolabeling. The relative immunolabeling efficiencies of the detecting antibodies were calculated based on the ratio of labeled proteins to the total number of proteins immobilized on the surface. Error bars represent means±s.d.

**Extended Data Fig. 6.**
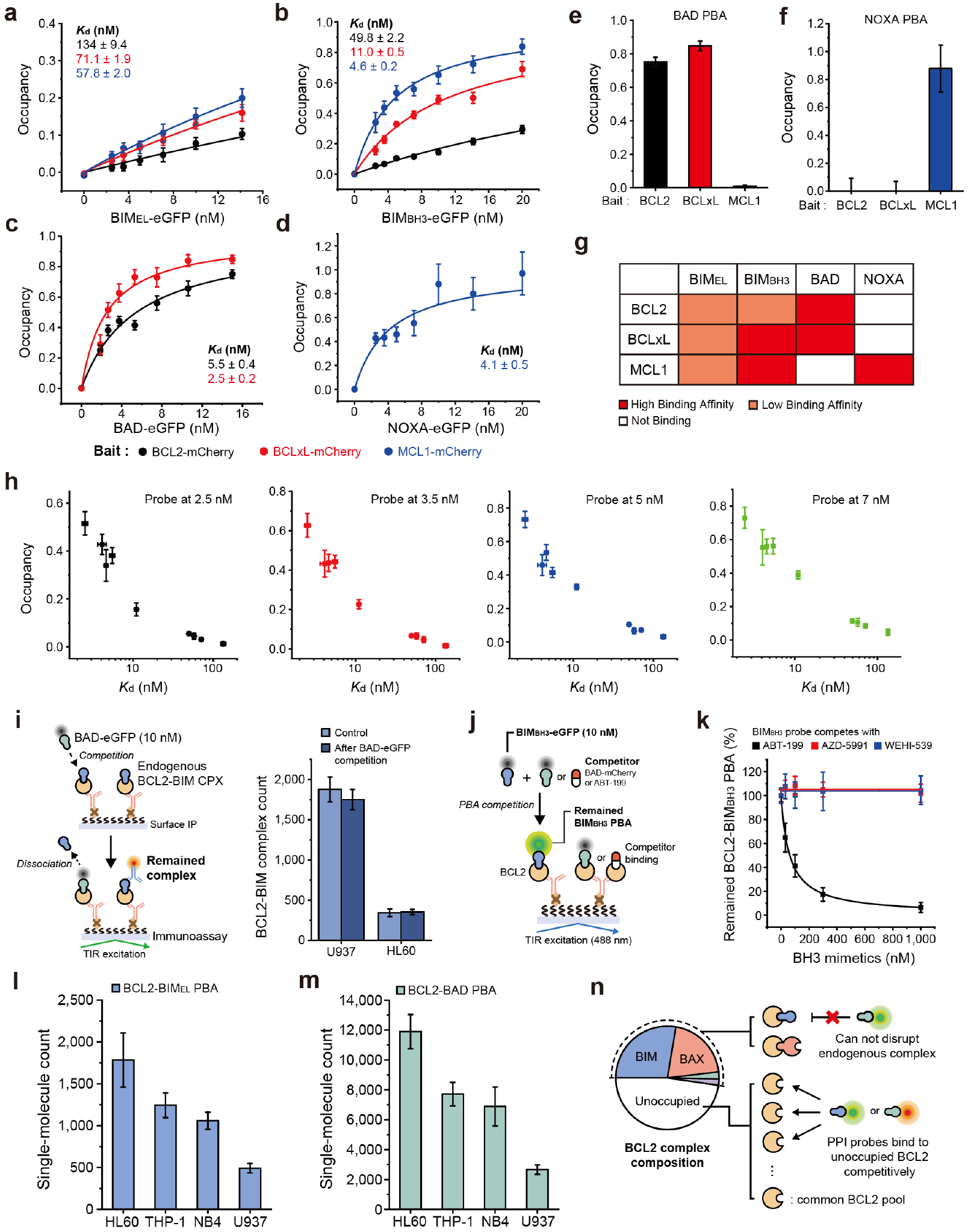
| Population profile of PPI probes binding onto unoccupied BCL2 and the cartoon schematic of the experiment. **a-d**, PBA binding curves for each PPI probes (eGFP-labeled BIM_EL_, BIM_BH3_, BAD, NOXA) to anti-apoptotic proteins (mCherry-labeled BCL2, BCLxL, MCL1) to calculate dissociation constant (*K*_d_). (**a**) BIM_EL_-PBA, (**b**) BIM_BH3_-PBA, (**c**) BAD-PBA, (**d**) NOXA-PBA. **e,f,** PBA binding patterns of each PPI probe. (**e**) BAD-PBA, (**f**) NOXA-PBA. **g**, Comparison of the binding affinities between each binding pairs. **h,** Correlations between the calculated *K*_d_ values and the occupancy values at a fixed PPI probe concentration. **i,** Dissociation of BCL2-BIM complex after *in vitro* competition by BAD-eGFP PPI probes. Protein complexes were surface-immobilized by anti-BCL2 IP antibody and detected with anti-BIM detection antibody after the competition (inset). **j,** Schematic of *in vitro* PBA competition between BIM_BH3_-eGFP PPI probe and competitors (PPI probe or BH3 mimetics) on surface-immobilized BCL2 proteins. **k**, Remained BCL2-BIM_BH3_ PBA after *in vitro* binding competition with BH3 mimetics (ABT-199, AZD-5991, WEHI-539). BIM_BH3_ PPI probe was presented in 10 nM. **l,m,** BCL2 PBA counts with different PPI probes from four AML cell lines (HL60, THP-1, NB4, U937). (**l**) BCL2-BIM_EL_ PBA, (**m**) BCL2-BAD PBA. **n**, Schematic for selective binding of PPI probes for unoccupied BCL2 proteins. Error bars represent means±s.d.

**Extended Data Fig. 7.**
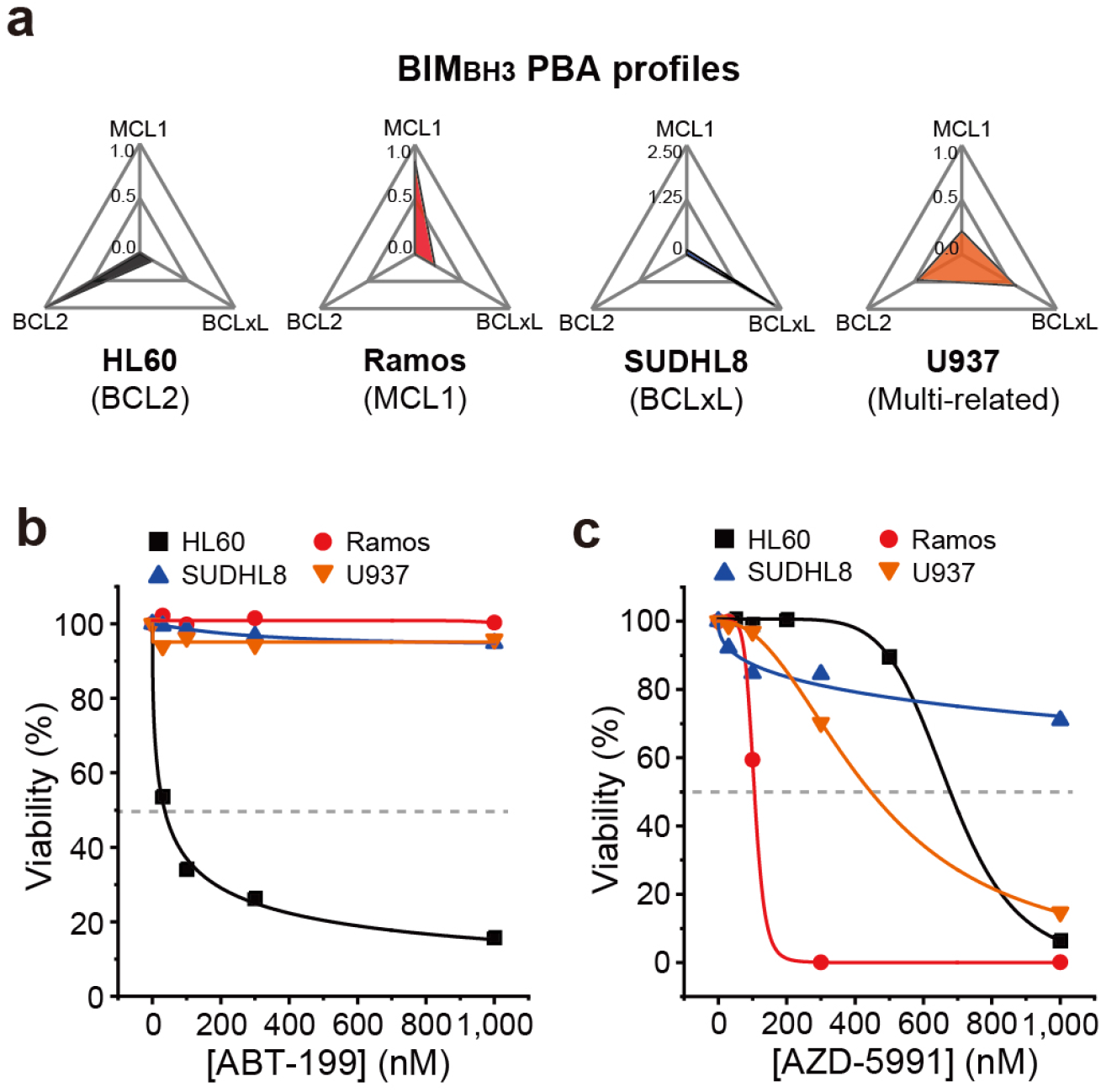
| BIM_BH3_ PBA profile predicts response of single-agent BH3 mimetics on leukemia cell lines. **a**, BIM_BH3_ PBA profiles of four different leukemia cell lines (HL60, Ramos, SUDHL8, U937). The BIM_BH3_ PBA levels were rescaled to account for the relative binding affinities of BIM_BH3_ probe for anti-apoptotic proteins determined in Extended Data Fig .6b. **b,c**, Viability of leukemia cell lines after treatment of BH3 mimetics for 24 hours. Viability represents the double negative populations after Annexin V/PI staining and analyzed with flow cytometry. (**b**) ABT-199, (**c**) AZD-5991.

**Extended Data Fig. 8.**
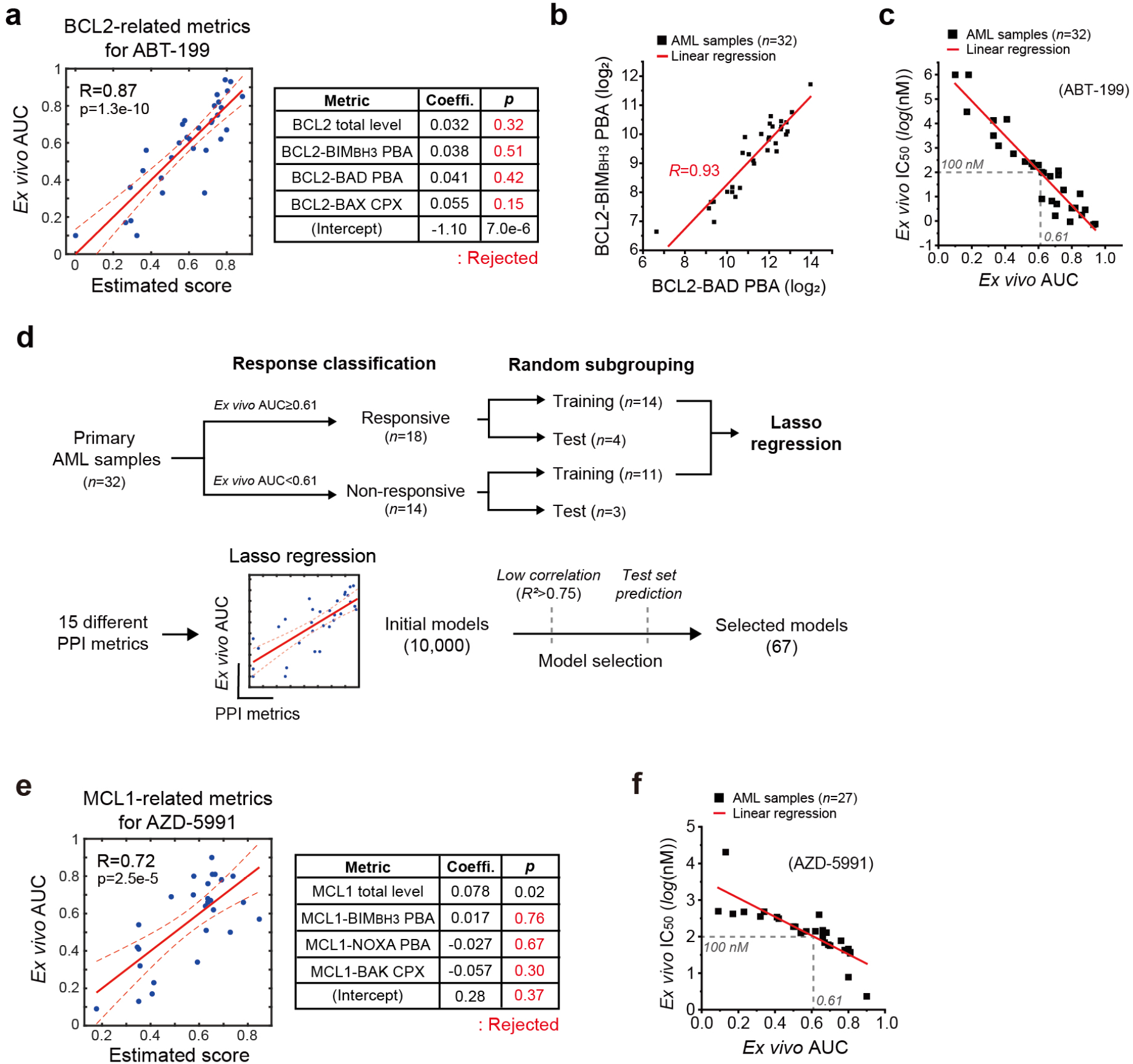
| Development of the *ex vivo* drug efficacy prediction model with combination of multiple-PPI metrics. **a**, Correlation between *ex vivo* AUC for ABT-199 and combination of BCL2-related metrics (BCL2 total level, BCL2-BIM_BH3_ PBA, BCL2-BAD PBA, and BCL2-BAX CPX). **b,** Linear correlations between two different BCL2 PBA metrics. **c,** Correlations between *ex vivo* AUC and IC_50_ of primary AML samples for ABT-199 (*n*=32). **d,** Schematic of Lasso regression analysis for the selection of PPI metrics highly correlated with drug response. The training and test groups were randomly selected from the primary AML sample cohort. The Lasso regression models were generated using the training group and evaluated based on Pearson’s R as well as prediction outcomes for the test group. 67 models were selected from 10,000 different initial models. **e,** Correlation between *ex vivo* AUC for AZD-5991 and combination of MCL1-related metrics (MCL1 total level, MCL1-BIM_BH3_ PBA, MCL1-NOXA PBA, and MCL1-BAK CPX). **f,** Correlations between *ex vivo* AUC and IC_50_ of primary AML samples for AZD-5991 (*n*=27).

**Extended Data Fig. 9.**
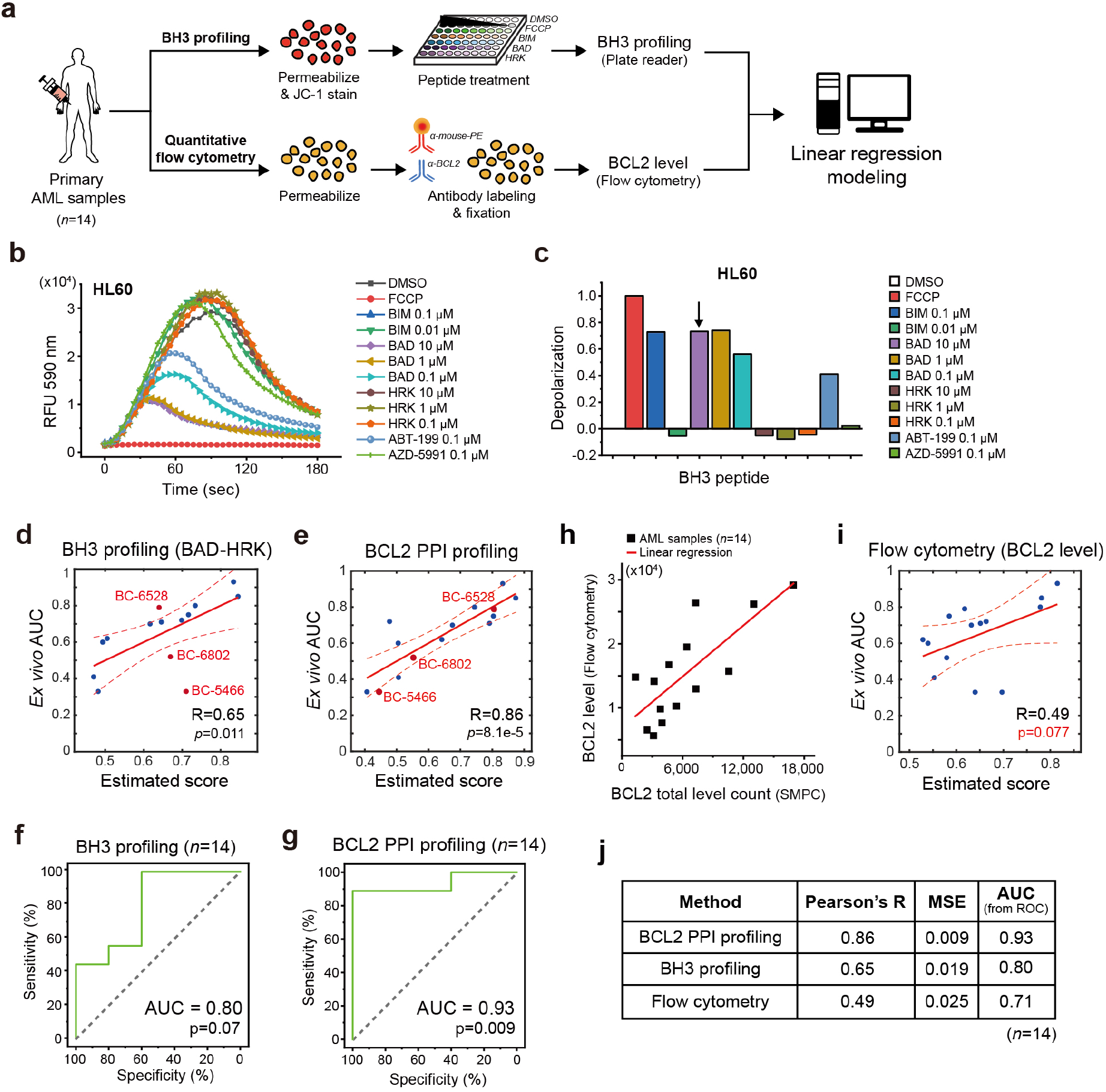
| Comparison of BCL2 PPI profiling and different methods to predict BH3 mimetics response. **a**, Schematic for BH3 profiling based on JC-1 staining (upper) and measurement of BCL2 levels using flow cytometry (lower) for primary AML samples (*n*=14). **b,c,** BH3 profiling on HL60 cells. (**b**) Relative fluorescence units (RFU) of 590 nm wavelength for HL60 cells through the treatment of BH3 peptides (BIM, BAD, and HRK) or BH3 mimetics (ABT-199 and AZD-5991). (**c**) Depolarization of HL60 cells through the BH3 profiling. Depolarization by 10 μM of BAD peptides was indicated. **d,e,** Correlations for *ex vivo* ABT-199 AUC with (**d**) depolarizations by BAD (10 μM) – HRK (10 μM) peptides, (**e**) combination of multiple PPI metrics (BCL2-BIM_BH3_ PBA, BCL2-BAX CPX, and BCLxL-BAK CPX). The outliers identified in model (**d**) were indicated. **f,g,** Receiver operating characteristic (ROC) curve for *ex vivo* ABT-199 responses with (**f**) depolarizations by BAD (10 μM) – HRK (10 μM) peptides, (**g**) Combination of multiple PPI metrics. **h,** Correlations between BCL2 protein levels determined by SMPC and flow cytometry for primary AML samples (n=*14*). **i,** Prediction models for ABT-199 *ex vivo* AUC with BCL2 protein levels determined by flow cytometry. **j,** Comparison of the predictive powers for ABT-199 drug responses across different methods (*n*=14).

**Extended Data Fig. 10.**
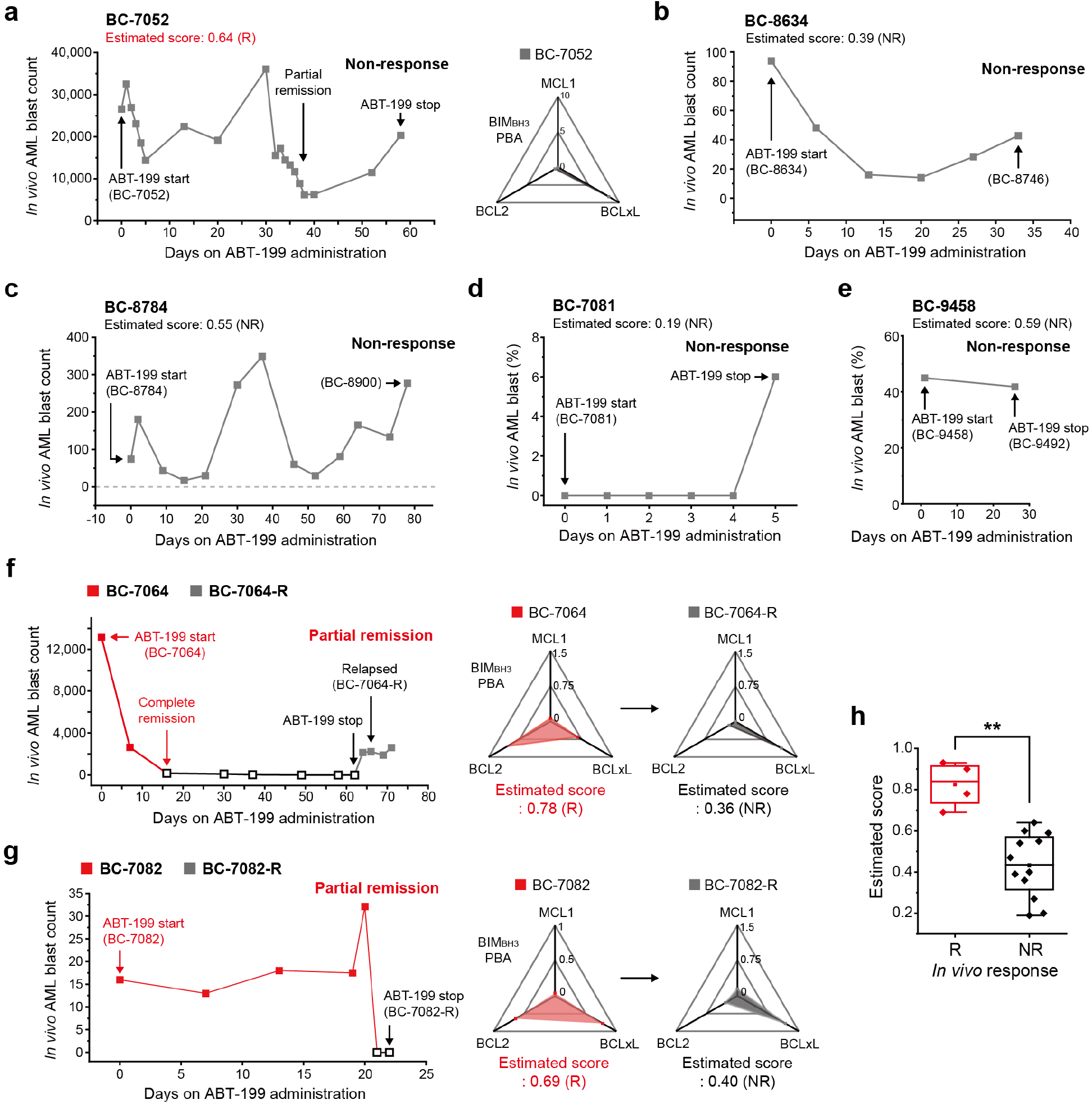
| Longitudinal tracking of *in vivo* ABT-199 response for AML patients. **a-g**, Changes of *in vivo* AML blast counts (or AML blast ratio) through the days after ABT-199 administration. (**a**) BC-7052, (**b**) BC-8634, (**c**) BC-8784, (**d**) BC-7081, (**e**) BC-9458, (**f**) BC-7064, (**g**) BC-7082. (**a-g**) The estimated scores and the PPI diagnostic results (R: Responsive, NR: Non-responsive) were obtained from the model in Figure 4f. The BIM_BH3_ PBA profiles of each sample were presented. **h**, Comparison of the estimated scores of AML samples between responsive (R, *n*=4) and non-responsive (NR, *n*=10) samples for *in vivo* ABT-199 administration (Two-sided Mann-Whitney test, *p*-value=0.004). Six additional samples from the same cohort collected after relapse (BC-7064-R, BC-7082-R, BC-7107-R2, BC-8900, BC-9458, BC-9492) were included, which were categorized as the NR.

